# Mapping and characterization of structural variation in 17,795 deeply sequenced human genomes

**DOI:** 10.1101/508515

**Authors:** Haley J. Abel, David E. Larson, Colby Chiang, Indraniel Das, Krishna L. Kanchi, Ryan M. Layer, Benjamin M. Neale, William J. Salerno, Catherine Reeves, Steven Buyske, NHGRI Centers for Common Disease Genomics, Tara C. Matise, Donna M. Muzny, Michael C. Zody, Eric S. Lander, Susan K. Dutcher, Nathan O. Stitziel, Ira M. Hall

## Abstract

A key goal of whole genome sequencing (WGS) for human genetics studies is to interrogate all forms of variation, including single nucleotide variants (SNV), small insertion/deletion (indel) variants and structural variants (SV). However, tools and resources for the study of SV have lagged behind those for smaller variants. Here, we used a cloud-based pipeline to map and characterize SV in 17,795 deeply sequenced human genomes from common disease trait mapping studies. We publicly release site-frequency information to create the largest WGS-based SV resource to date. On average, individuals carry 2.9 rare SVs that alter coding regions, which affect the dosage or structure of 4.2 genes and account for 4.0-11.2% of rare high-impact coding alleles. Based on a computational model, we estimate that SVs account for 17.2% of rare alleles genome-wide whose predicted deleterious effects are equivalent to loss-of-function (LoF) coding alleles; ~90% of such SVs are non-coding deletions (mean 19.1 per genome). We report 158,991 ultra-rare SVs and show that ~2% of individuals carry ultra-rare megabase-scale SVs, nearly half of which are balanced and/or complex rearrangements. Finally, we exploit this resource to infer the dosage sensitivity of genes and non-coding elements, revealing strong trends related to regulatory element class, conservation and cell-type specificity. This work will help guide SV analysis and interpretation in the era of WGS.

## INTRODUCTION

Human genetics studies employ WGS to enable comprehensive trait mapping analyses across the full diversity of genome variation, including SVs (≥50 bp) such as deletions, duplications, insertions, inversions and other rearrangements. Analyses of SV suggest that such variation plays a disproportionately large role (relative to their abundance) in rare disease biology^1^, and in shaping heritable gene expression differences in the human population^2–4^. Rare and *de novo* SVs have been implicated in the genetics of autism^5–8^ and schizophrenia^9–12^, but few other complex trait association studies have directly assessed SVs^13,14^. With the advent of WGS, it is now possible to characterize a more diverse collection of SVs and deepen our understanding across a wider range of traits.

One challenge for SV interpretation in WGS-based studies is the relative lack of high-quality publicly available variant maps from large populations. Our current knowledge of SV in human populations is based primarily on three sources: (1) a large and disparate collection of array-based studies, organized in various databases^15–17^, with limited allele frequency information and technical challenges that limit sensitivity, accuracy and resolution; (2) the 1000 Genomes Project callset derived from 2,504 low-coverage (median 7x) WGS datasets^4^, which has been invaluable but is limited by the modest sample size and low coverage design (which hindered rare variant discovery); and (3) an assortment of smaller WGS-based studies with varied coverage, technologies, analysis methods, and levels of data accessibility, only 4 of which contribute more than 100 unrelated samples (500 Dutch^18^, 100 Danish^19^, 232 globally diverse^20^, and ~519 autism families^7,8^).

There is an opportunity to improve knowledge of SV in human populations via systematic analysis of large-scale WGS data resources generated by programs such as the NHGRI Centers for Common Disease Genomics (CCDG). A key barrier to the creation of larger and more informative SV catalogs is the lack of computational tools that can scale to the size of ever growing datasets. To begin to address this need, we have developed a highly scalable open source SV analysis pipeline (https://www.biorxiv.org/content/early/2018/12/13/494203), and used it to map and characterize SV in 17,795 deeply sequenced human genomes. Our results illuminate the landscape of rare SV in unprecedented detail. This SV catalog is freely available and will aid variant interpretation in "n-of-one" WGS applications.

## RESULTS

### A population-scale map of structural variation

The samples analyzed here are derived from common disease case/control and quantitative trait mapping collections sequenced under the CCDG program, supplemented with ancestrally diverse samples from the PAGE consortium (~950 Latinos) and Simons Genome Diversity Panel (263 individuals from 142 populations). The sample set also includes 1,823 families consisting of two or more first-degree relatives, including a set of multi-generational CEPH pedigrees comprising 595 samples and 80 founders. The samples do not include known rare disease cases. The final ancestry composition is extremely diverse, including 23% African American, 16% Latino, 11% Finnish European, 39% non-Finnish European, 6% unknown, and 6% ancestrally diverse samples from various sites around the world (Table 1).

**Table 1.**
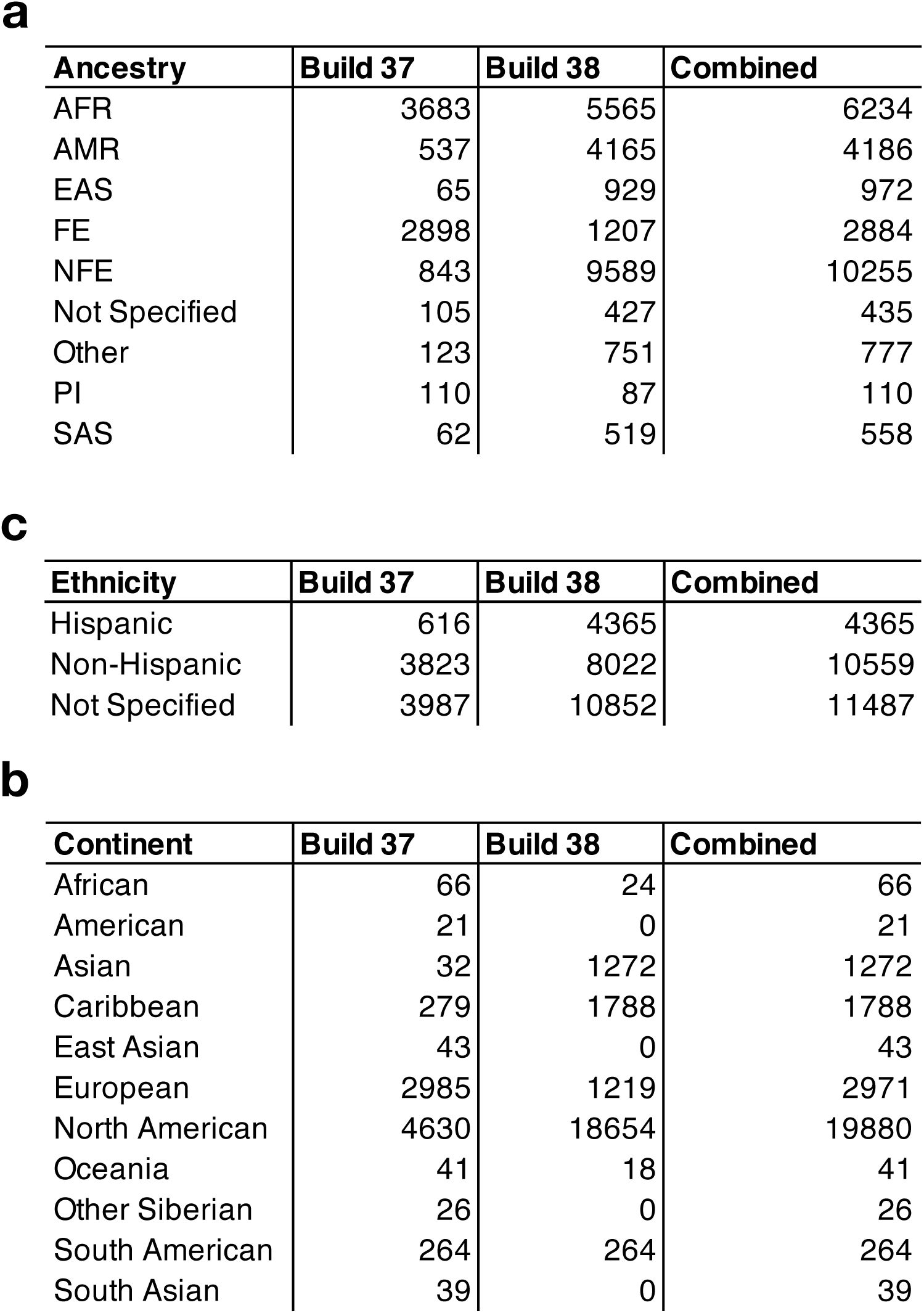
**(a)** Ancestry, **(b)** ethnicity, and **(c)** continental origin of the samples analyzed in this study. For each table, the number of samples in the B37 and B38 callsets are shown separately, including the non-redundant union at right. Abbreviations are as follows: AFR, African; AMR, admixed American; EAS, east Asian; FE, Finnish European; NFE, non-Finnish European; PI, Pacific Islander; SAS, South Asian.

| Ancestry | Build 37 | Build 38 | Combined |
| --- | --- | --- | --- |
| AFR | 3683 | 5565 | 6234 |
| AMR | 537 | 4165 | 4186 |
| EAS | 65 | 929 | 972 |
| FE | 2898 | 1207 | 2884 |
| NFE | 843 | 9589 | 10255 |
| Not Specified | 105 | 427 | 435 |
| Other | 123 | 751 | 777 |
| PI | 110 | 87 | 110 |
| SAS | 62 | 519 | 558 |

**c**
| Ethnicity | Build 37 | Build 38 | Combined |
| --- | --- | --- | --- |
| Hispanic | 616 | 4365 | 4365 |
| Non-Hispanic | 3823 | 8022 | 10559 |
| Not Specified | 3987 | 10852 | 11487 |

**Table 1. (a)** Ancestry, **(b)** ethnicity, and **(c)** continental origin of the samples analyzed in this study.
| Continent | Build 37 | Build 38 | Combined |
| --- | --- | --- | --- |
| African | 66 | 24 | 66 |
| American | 21 | 0 | 21 |
| Asian | 32 | 1272 | 1272 |
| Caribbean | 279 | 1788 | 1788 |
| East Asian | 43 | 0 | 43 |
| European | 2985 | 1219 | 2971 |
| North American | 4630 | 18654 | 19880 |
| Oceania | 41 | 18 | 41 |
| Other Siberian | 26 | 0 | 26 |
| South American | 264 | 264 | 264 |
| South Asian | 39 | 0 | 39 |

The tools and pipelines used for this work are described in detail elsewhere (https://www.biorxiv.org/content/early/2018/12/13/494203). Briefly, we have developed a highly scalable software toolkit (svtools) and distributed workflow for large-scale SV callset generation that combines per-sample variant discovery (via LUMPY^21^), resolution-aware cross-sample merging, breakpoint genotyping (via SVTYPER^22^), copy number annotation, variant classification, and callset refinement (Fig. 1). Our pipeline is built on a foundation of well-established tools (LUMPY^21^, SPEEDSEQ^22^, SVTYPER^22^) that have been extensively tested in prior studies^3,21,22^, combined with various new optimizations and an innovative cross-sample merging and genotyping strategy that provides efficient and affordable joint analysis of >100,000 genomes at near-base-pair resolution.

**Figure 1.**
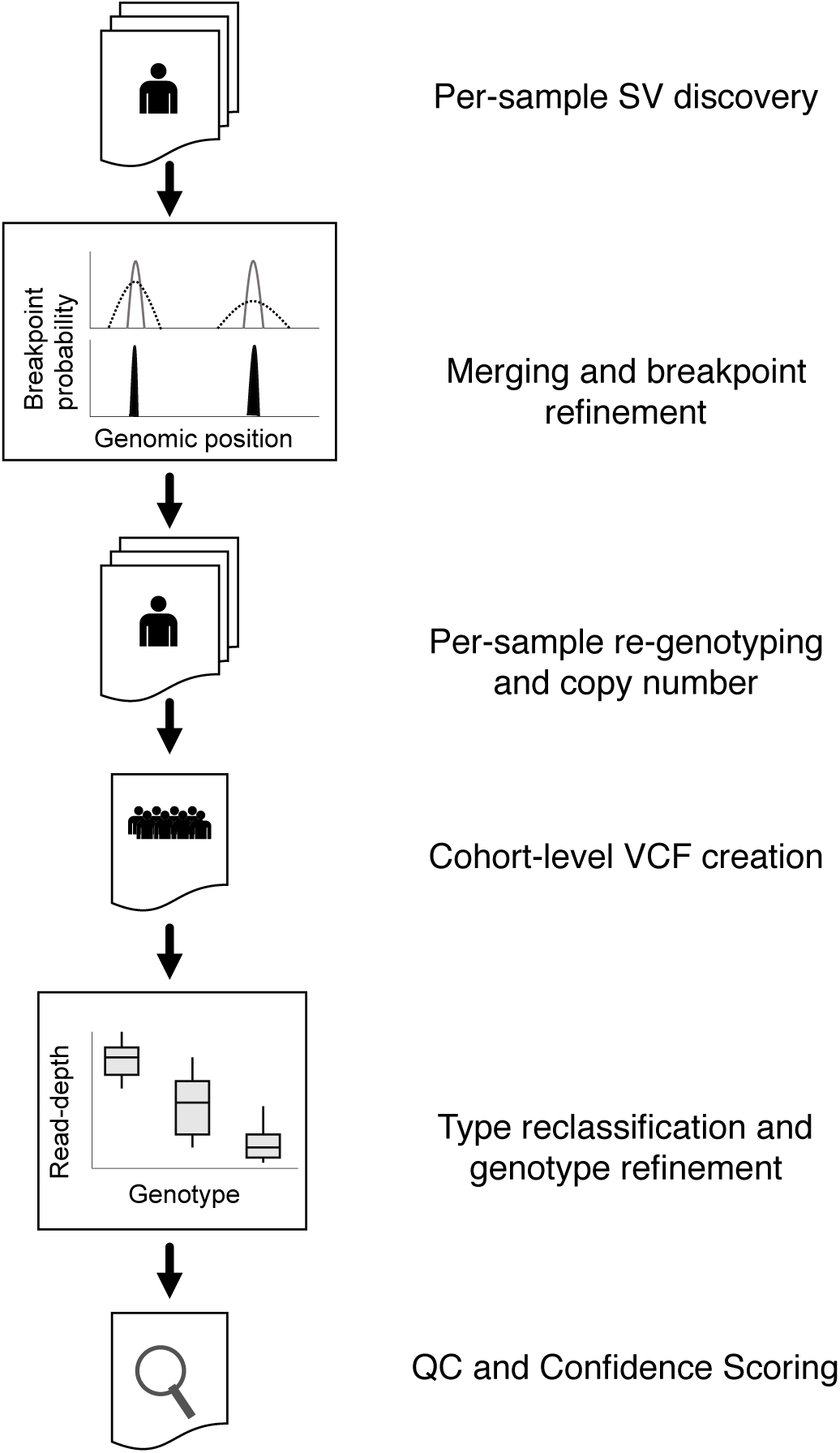
Callset construction pipeline. SV are detected within each sample using LUMPY. Breakpoint probability distributions are used to merge and refine the position of detected SV within a cohort, followed by parallelized re-genotyping, and copy number annotation. Samples are merged into a single, cohort-level VCF file, variant types reclassified and genotypes refined with svtools using the combined breakpoint genotype and read-depth information. Finally, sample-level QC and variant confidence scoring is conducted to produce the final callset.

We created two distinct SV callsets using different reference genome and pipeline versions. The "B37" callset includes 118,973 high-confidence SVs from 8,417 samples sequenced at the McDonnell Genome Institute, that were aligned and processed using SpeedSeq^22^ and the GRCh37 reference genome and analyzed with the LSF-based "B37" SV mapping pipeline. The "B38" callset includes 241,426 high-confidence SVs from 23,239 samples sequenced at four different CCDG sites, that were aligned to GRCh38 using the newly-developed “functional equivalence” pipelines^23^, and analyzed on the Google Cloud using the "B38" SV mapping pipeline (see **Methods**) as part of CCDG "Freeze 1". 5,245 samples are included in both callsets, resulting in a non-redundant total of 26,731 samples. Of these, 17,795 samples are permitted for aggregate-level sharing; these comprise the official public release (**Supplementary Files 1 and 2**) and are the basis for all analyses presented below. This dataset contains ~7-fold more individuals than the largest prior WGS-based study of SV^4^. We release the public B37 (n=8,417) and B38 (n=14,623) callsets separately to enable their use in applications that are sensitive to reference genome and pipeline version effects, although they can be combined (e.g., via liftover) for basic analyses involving unique genomic regions, as below. For simplicity, most analyses below focus on the public version of the B38 callset, although we note that the B37 callset will be extremely useful for the numerous human genetics projects that continue to rely on the GRCh37 reference.

By a variety of measures, both callsets appear to be high quality. We observe a mean of 4,442 high-confidence SVs per genome, predominantly composed of deletions (35%), mobile element insertions (27%), and tandem duplications (11%) (Fig. 2b, Supplementary Figs. 1 and 2). Variant counts and linkage disequilibrium patterns (Supplementary Figs. 1c and 1e) are consistent with prior studies employing similar methods, including the GTEx callset that was characterized extensively in the context of eQTL mapping^3^. We achieve expected variant detection performance for embedded 1KGP samples, consistent with prior studies using similar methods^3,21,22^ (https://www.biorxiv.org/content/early/2018/12/13/494203). As expected, the site-frequency spectrum largely mirrors that of SNVs and indels (Fig. 2c, Supplementary Fig. 1d), the size distribution shows increasing length with decreasing frequency (which suggests negative selection against larger variants, Fig. 2d), principal components analysis reveals the expected population structure based on self-reported ancestry (Supplementary Fig. 3), and dosage sensitivity analyses show strong concordance with independent measures of functional constraint and genome function (see Fig. 5 and analysis below). The number and types of high-confidence SVs observed per genome are remarkably consistent across the sample set (Supplementary Figs. 1 and 2), with noticeably higher levels of genetic variation in African-ancestry individuals, as expected. Although there is some technical variability in SV numbers due to cohort and sequencing center, the effects are mainly limited to small (<1 kb) deletions and tandem duplications detected solely by read-pair signals, which are sensitive to library preparation and alignment filtering methods that differ among CCDG sites (Supplementary Fig. 2a, see **Methods**). In this respect the B37 callset has superior quality due to the more consistent read-depth and fragment length distribution of WGS data produced at a single center. Finally, and perhaps most importantly, high-confidence SVs have an acceptably low Mendelian error rate (<5%; see Supplementary Figs. 1c **and** 2c) as judged by segregation within 36 nuclear families, and there is a strong relationship between variant quality metrics and error rate (Supplementary Fig. 2c), such that the callset can be further tuned for specific applications using available metadata.

**Figure 2.**
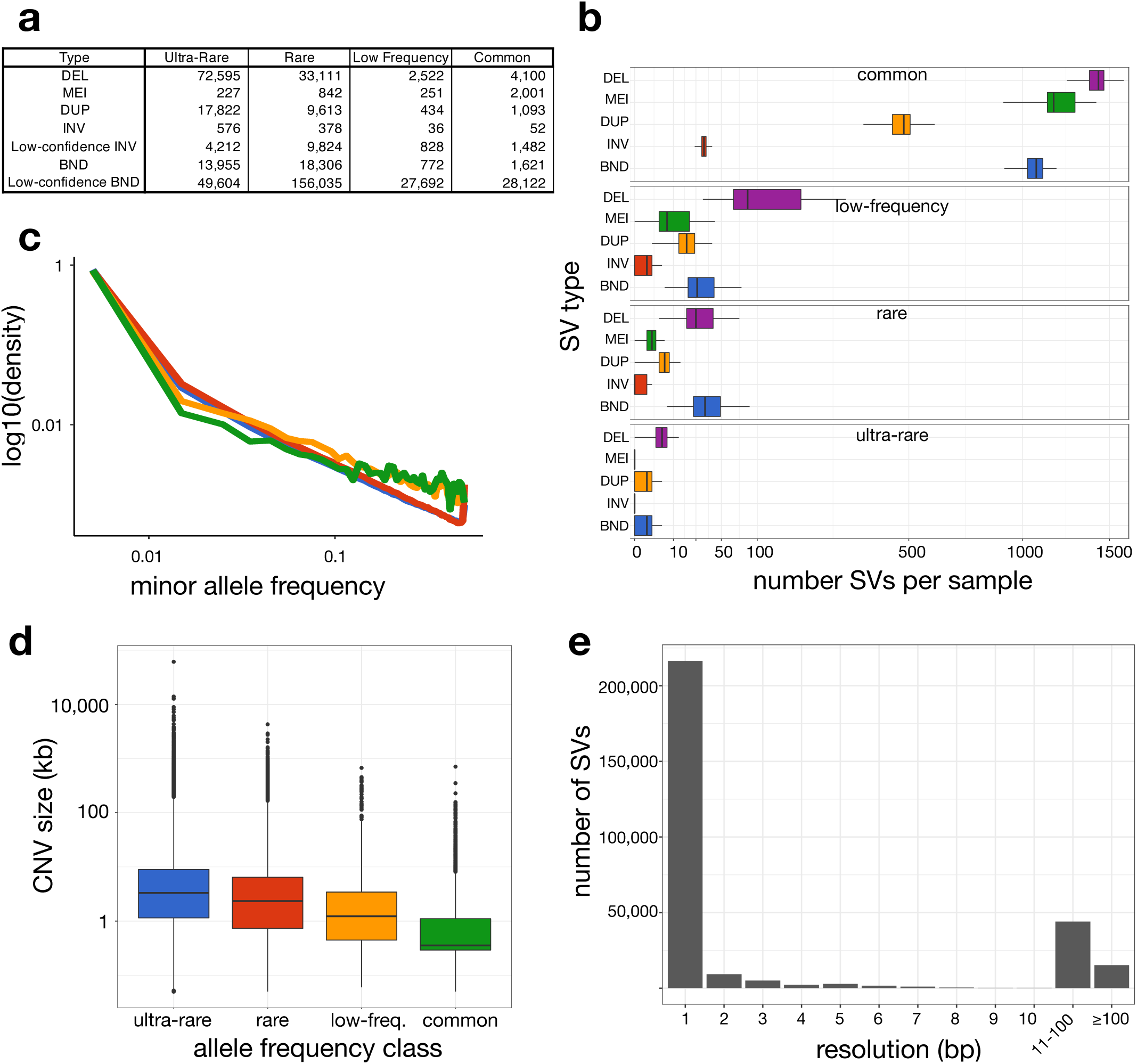
The public version of the “B38” callset derived from 14,623 samples. **(a)** Number of high-confidence and low-confidence SVs by class and frequency bin. SV classes are defined as: DEL, deletion; MEI, mobile element insertion; DUP, duplication; INV, inversion; BND, “break-end”, which is a generic term in the VCF specification for SV breakpoints that cannot be unequivocally classified. Minor allele frequency (MAF) bins are defined as: “ultra-rare” is private to an individual or family; “rare” is MAF<1%; “low-frequency” is 1%<MAF<5%; “common” is MAF>5%. **(b)** Number of SVs per sample (x-axis, square-root scaled) by SV type (y-axis) and frequency class (panels labelled at top). **(c)** MAF distribution for SNV, indel, deletion (DEL) and duplication (DUP) variants for a subset of 4,298 samples for which GATK-based SNV/indel were also available. **(d)** CNV length distributions for each frequency class, defined as in part (a). **(e)** Histogram showing the resolution of SV breakpoint calls, as defined by the length of the 95% confidence interval of the breakpoint-containing region defined by LUMPY, after cross-sample merging and refinement using svtools.

Notably, both SV maps have extremely high genomic resolution relative to current resources: 72% of SV breakpoints are mapped to single-base resolution and 80% are mapped within 10 bp (Fig. 2e). The high resolution nature of this resource will be extremely valuable for variant interpretation in n-of-one studies, and for developing graph-based SV genotyping methods^24^.

### Burden of deleterious rare structural variants

The contribution of rare SV to human disease remains unclear. Well-powered WGS-based trait mapping studies will ultimately be required to address this; however, the overall burden of predicted pathogenic mutations in the human population is informative and can be estimated from our data. Our joint analysis of 14,623 individuals (B38 callset) identified 42,765 rare SV alleles (MAF<1%) predicted to decrease gene dosage (n=9,416), alter gene function (e.g., single exon deletion; n=26,337), or increase gene dosage (n=7,012). The majority of rare gene-altering SVs are deletions (54.5%), with somewhat fewer duplications (42.2%), and a small fraction of other variant types (3.3%) primarily composed of inversions and complex rearrangements that interrupt or rearrange exons. Of these, 23.4% affect multiple genes and 10.4% affect 3 or more genes, resulting in a mean of 4.2 SV-altered genes per individual. If we define SV-based loss-of-function (LoF) mutations strictly to encompass gene disruptions and gene deletions affecting >20% of exons, we identify a mean of 1.39 rare SV-based gene LoF alleles per person. Analysis of a subset of 4,298 individuals with both SV and SNV/indel calls reveals that individuals carry a mean of ~33.6 rare high-confidence LoF mutations caused by SNVs or small indels (Fig. 3a) which is consistent with prior studies^25^. This result shows that SVs account for 4.0-11.2% of rare, predicted high impact gene alterations in a population sample of individuals, depending on whether we consider all coding SVs, or a strictly defined set of LoF variants. These are likely to be underestimates considering that the false negative rate of SV detection is typically higher than that of SNVs and small indels^22^ (https://www.biorxiv.org/content/early/2018/06/13/193144).

**Figure 3.**
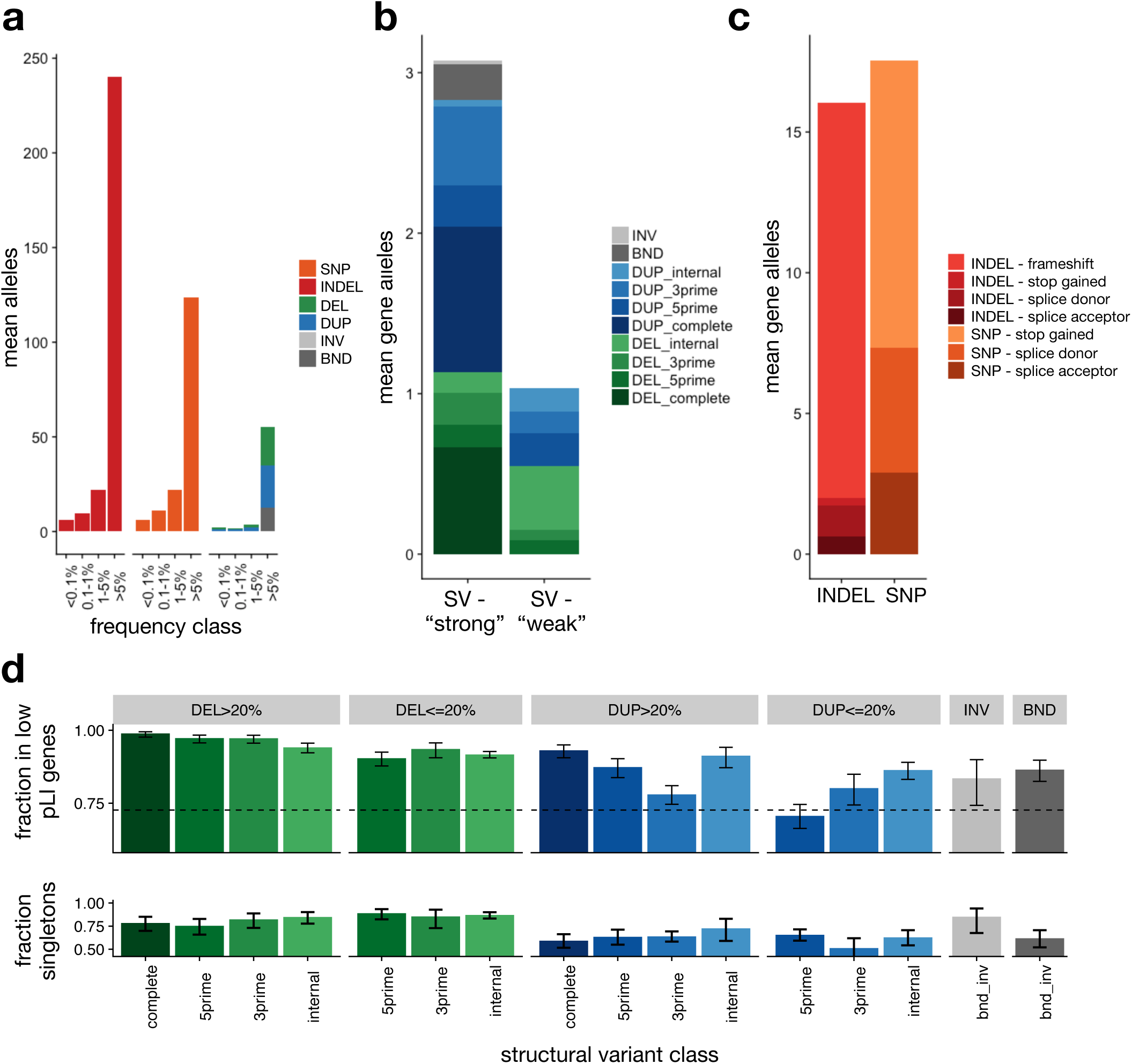
Burden of rare gene-altering SV. **(a)** Per-sample mean number of gene-altering variants by type and frequency class. **(b)** Per-sample mean number of rare (<1% MAF) gene-altering SV by type. DEL and DUP are subclassified into ‘strong’ (affecting >20% of exons of principal transcript) and ‘weak’ (affecting <20% of exons of principal transcript and ‘internal’ (variant overlaps at least one coding exon, but neither the 3’ nor 5’ end of the principal transcript), 3prime (variants overlaps the 3’ end of the transcript), 5prime (variant overlap the 5’ end of the transcript), and complete (variant overlaps all coding exons in principal transcript). **(c)** Per-sample mean number of rare (<1% MAF) high-confidence PTV by type and vep consequence. **(d)** (top) Fraction of rare (<1% MAF), gene-altering variants occurring in low pLI (pLI<0.9) vs. high pLI (pLI>=0.9) genes, by type, size class, and gene region. Error bars indicate 95% confidence intervals (Wilson score interval). The dotted line indicates the expected fraction, assuming a uniform distribution of SV in coding exons. (bottom) Singleton rates for gene-altering variants by type, restricted to genes with pLI>0.1. Error bars indicate 95% Wilson score confidence intervals.

To characterize the relative impact of different coding SV classes we calculated two measures of purifying selection (Fig. 3d): (1) the fraction of variants that affect dosage tolerant genes with an LoF intolerance (pLI)^25,26^ score <0.9, which reflects depletion of that variant class in more dosage sensitive genes; and (2) the fraction of variants present as ultra-rare "singletons" found in only one individual or family, which reflects the extent to which alleles of a given class are being "flushed" from the population due to their deleterious effects (under the assumptions outlined below). Deletions are more deleterious than duplications, complete gene deletions are the most deleterious class, and sub-genic deletions affecting ≥20% of exons approach similar levels as whole gene deletions. Notably, based on the fraction of variants in dosage tolerant genes, complete gene duplications and sub-genic deletions affecting <20% of exons show surprisingly strong levels of deleteriousness. This suggests that most gene-altering SVs are strongly deleterious, even if not predicted to completely obliterate gene function, and that the upper range of our 4.0-11.2% estimate may be more accurate.

The above calculations ignore deleterious missense and non-coding variants that are expected to comprise a large fraction of rare functional variation. Predicting the impact of these variant types is challenging; however, it should be possible to approximate their proportional contribution to the deleterious variant burden under two simplifying assumptions: (1) impact prediction algorithms such as CADD^27^ and LINSIGHT^28^ are capable of ranking variants within a given class (SNV, indel, SV) by their degree of deleteriousness, and (2) the mean deleteriousness of a given subset of variants is reflected by its singleton rate. The first assumption is somewhat tenuous, but should be valid for this analysis considering that impact prediction inaccuracies are likely to affect variant classes similarly given use of the same underlying models (CADD and LINSIGHT). The second assumption should hold true under an infinite sites model of mutation, which seems reasonable at the sample size used for this analysis (n=4,298). We note that other evolutionary forces such as positive selection, background selection, and biased gene conversion can also shape the site frequency spectrum; however, it seems likely that these forces would act similarly on the variant classes examined here, in a genome-wide analysis of a very large number of sites.

Thus, to compare levels of deleterious SVs relative to SNVs and indels, we can select impact score thresholds separately to yield the same singleton rate for each class. We used CADD and LINSIGHT to generate combined impact scores for SNVs, indels, deletions and duplications (see **Methods**), where SVs are scored using a scheme that aggregates per-based CADD/LINSIGHT scores across the affected genomic region (as in SVScore^29^). As expected, these scores are highly correlated with singleton rate and with variant effect predictions from VEP^30^ and LOFTEE^25^ (Fig. 4a-c). We sought to identify variants from each class (SNV, indel, DEL, and DUP) with deleteriousness equivalent to high-confidence LoF mutations defined by VEP and LOFTEE, by using an impact score threshold that yields a singleton rate matching that of the entire set of high-confidence LoF mutations (hereafter referred to as "strongly deleterious" variants). Individuals carried a mean of 121.9 such “strongly deleterious” rare variants, comprising 63% SNVs, 19.8% indels and 17.2% SVs (96.9% of which are deletions) (Fig. 4d). Taking into account the relative numerical abundance of different rare variant classes, this suggests that a given rare SV is 841-fold more likely to be strongly deleterious than a rare SNV, and 341-fold more likely than a rare indel. Predicted deleterious SVs are slightly larger than rare SVs on the whole (median 4.5 vs. 2.8 kb). Whereas only a minority (13.1%) of predicted strongly deleterious SNVs and indels are non-coding, 90.1% of predicted strongly deleterious rare SVs are non-coding. Remarkably, the top 50% of non-coding deletions show similar levels of purifying selection (as measured by singleton rate) as high-confidence LoFs caused by SNVs/indels (see Fig. 4c), implying that a typical individual carries 19.1 strongly deleterious rare non-coding deletion alleles. This suggests that non-coding deletions have surprisingly strong deleterious effects and may play a larger than expected role in human disease.

**Figure 4.**
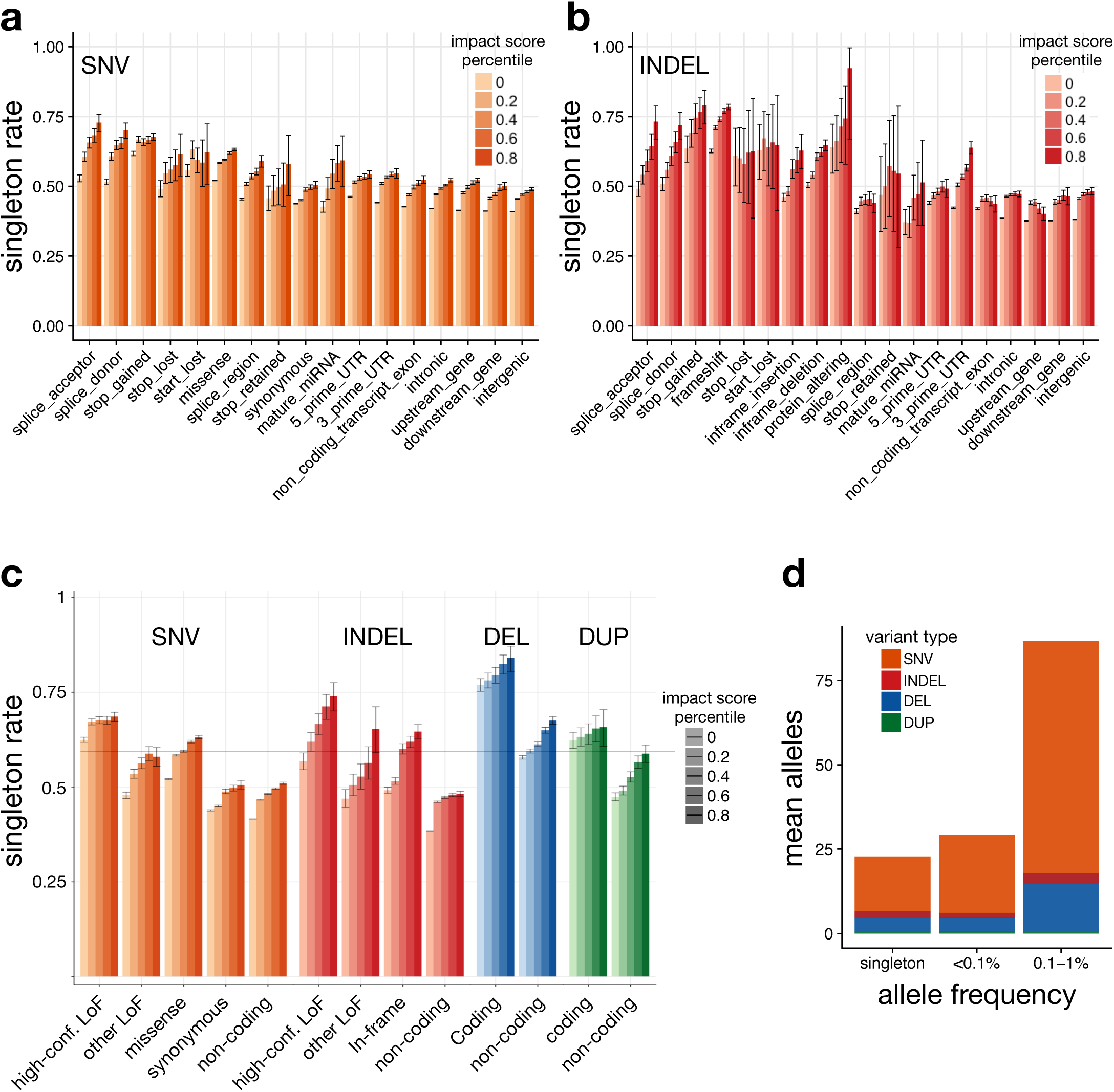
Estimation of genome-wide burden of functional alleles. **(a)** Singleton rates for SNV, by VEP consequence and percentile of combined VEP/CADD impact score. **(b)** Singleton rates for indels. **(c)** Singleton rates by variant type and percentile of combined VEP/CADD impact score. Here, ‘other LoF’ indicates VEP-annotated protein-truncating variants (PTVs) that are not classified as high-confidence by LOFTEE. DELs and DUPs that intersect any coding exon of the principal transcript are classified as ‘coding’; otherwise they are ‘noncoding’. **(d)** Per-sample mean number of ‘high’ impact alleles genome-wide, by type and frequency class.

These results demonstrate that SVs comprise a significant fraction of the burden of rare deleterious variants in the human population, to an extent that greatly exceeds their numerical contribution to genome variation overall, and that comprehensive ascertainment of SVs will improve trait association power in both common and rare disease studies.

### Landscape of ultra-rare structural variation

Most ultra-rare SVs represent recent or *de novo* structural mutations, and thus the relative abundance of different ultra-rare SV classes sheds light on the underlying mutational processes at work. We identified 158,991 ultra-rare SVs (105,175 high-confidence) that were present in only one of the 14,623 individuals included in the B38 callset, or that were private to a single family (MAF<0.01%). This corresponds to a mean number of ~11.4 per individual, with a relatively uniform distribution across individuals (Supplementary Fig. 4a). Ultra-rare SVs are mainly composed of deletions (5.2 per person) and duplications (1.3), with a smaller number of inversions (0.17). Interestingly, ~40% of ultra-rare SV breakpoints in our dataset cannot be readily classified into the canonical forms of SV. This is a known limitation of short-read WGS, and often such variants are ignored. Formally, as per the VCF specification^31^, these SVs are of the generic "breakend" (BND) variant class used to denote SVs whose true structure is unknown. We examined the 63,559 ultra-rare BNDs for insights into their composition and origin. Many (17.0%) appear to be deletions that are simply too small (<100 bp) to exhibit convincing read-depth support, and that our pipeline conservatively classifies as BNDs rather than assuming DNA has been lost (e.g., complex SVs can masquerade as deletions). 2.4% of the ultra-rare BNDs stem from 1,542 "retrogene insertions" caused by retroelement machinery acting on mRNAs. This map of retrogene insertions (also known as "retrocopies" or "retroduplications") is ~10-fold larger than prior maps^32–34^ and will be valuable for future studies of this phenomenon. 5.5% of ultra-rare BNDs appear to be complex genomic rearrangements with multiple breakpoints in close proximity (<100 kb). The remainder are difficult-to-classify variants involving either local (49.9% ≤1 Mb), distant intra-chromosomal (5.7% >1 Mb), or inter-chromosomal alterations (27.2%), many (78.0%) of which are classified as low-confidence SV calls. This final class is likely caused primarily by repetitive element variation, but should be interpreted with caution since we also expect them to be enriched for false positives.

A variety of sporadic disorders are caused by extremely large and/or complex SVs, but the frequency of these dramatic alterations in the general population is not well understood. These include megabase-scale CNVs, translocations, and complex genomic rearrangements involving multiple distant loci. We observed 47 extremely large (>1 Mb) deletions and 91 extremely large duplications, corresponding to a frequency of ~0.01 per individual, which affect a mean of 12.1 genes (Supplementary Fig. 4b). Three individuals carried two megabase-scale CNVs, apparently due to independent mutations. We observed 19 reciprocal translocations, corresponding to a frequency of 0.001 per individual, consistent with (albeit somewhat lower than) prior cytogenetic-based estimates^35,36^. Of these translocations, 14 affect a gene at one breakpoint and 2 affect a gene at both breakpoints, producing 1 predicted in-frame gene fusions (PI4KA:MGLL). We next applied breakpoint clustering approaches (as in^37^) to identify ultra-rare complex rearrangements involving 3 or more breakpoints and discovered 33 complex SVs spanning >1 Mb, representing a frequency of 0.003 per individual. Most megabase-scale complex SVs (20/33, 60.6%) appear to involve three breakpoints; however, we observed 5 large-scale rearrangements with 5 or more breakpoints. Notably, when the entire SV size distribution is considered, 3.3% of ultra-rare SVs are complex variants based on the presence of multiple adjacent breakpoints in the same individual, consistent with previous smaller-scale studies^38–42^.

### Dosage sensitivity of genes and noncoding elements

A motivation for creating population-scale SV maps is to annotate genomic regions based on their tolerance to dosage changes and structural rearrangements. These annotations can reveal the genes and non-coding elements that are most important (or dispensable) for human development and viability, and thus help interpret rare variants in n-of-one studies. The pLI score from ExAC/gnomAD^25,26^ has proven invaluable for this purpose and is based on more samples than analyzed here; however, pLI does not measure the effects of increased dosage, or include non-coding elements. Other CNV-based dosage tolerance maps are based on microarray^43^ or exome sequencing data^44^, and thus have poor resolution and coverage of non-coding regions.

The analyses presented here focus on a non-redundant union of the two callsets including 17,795 samples. We first generated DEL and DUP "sensitivity" scores for each gene based on the frequency of CNVs observed in our callsets, following the general approach of Ruderfer et al.^44^ (see **Methods**). The resulting scores are significantly correlated with the DEL and DUP scores from ExAC^44^, and with the DECIPHER haploinsufficiency score^43^ (which includes data from ExAC) (Supplementary Figs. 5 **and** 6). Despite their relatively modest correlations with each other, all three measures are equally informative based on comparison to pLI, which was generated using SNVs and indels from ExAC, and thus uses an independent set of variants. A combined score from multiple datasets performs better than any single score, and will be useful for interpreting rare SVs (**Supplementary File 3**).

We next performed a genome-wide analysis by measuring the frequency of dosage alterations in 1 kb sliding windows across the entire genome (see Methods). Based on the density of CNVs, our current dataset is not large enough to predict dosage sensitive non-coding elements based on the absence of variation; however, we can investigate the relative sensitivity of different genomic features, in aggregate. As expected, genic regions are highly depleted for dosage alterations in a manner that correlates with gene pLI (Fig. 5a). In fact, dosage sensitive genes confound analysis of non-coding elements since they cast "shadows" that extend into neighboring regions. Indeed, we observe a strong depletion of CNV near coding exons that varies depending on the proximity to the nearest exon as well as pLI of the corresponding gene (Fig. 5a). We therefore estimated odds ratios for depletion of CNV in each functionally annotated region, stratified by binned distance and pLI of the nearest exons, using a Cochran-Mantel-Haenszel estimator. Because adjacent windows are not strictly independent observations – i.e., CNV or features may overlap adjacent windows and induce spatial correlations – we used a blocked bootstrap resampling procedure to estimate robust confidence intervals. The resulting dosage sensitivity scores strongly correlate with independent measures of selective constraint including LINSIGHT and PHASTCONS (Fig. 5b).

**Figure 5.**
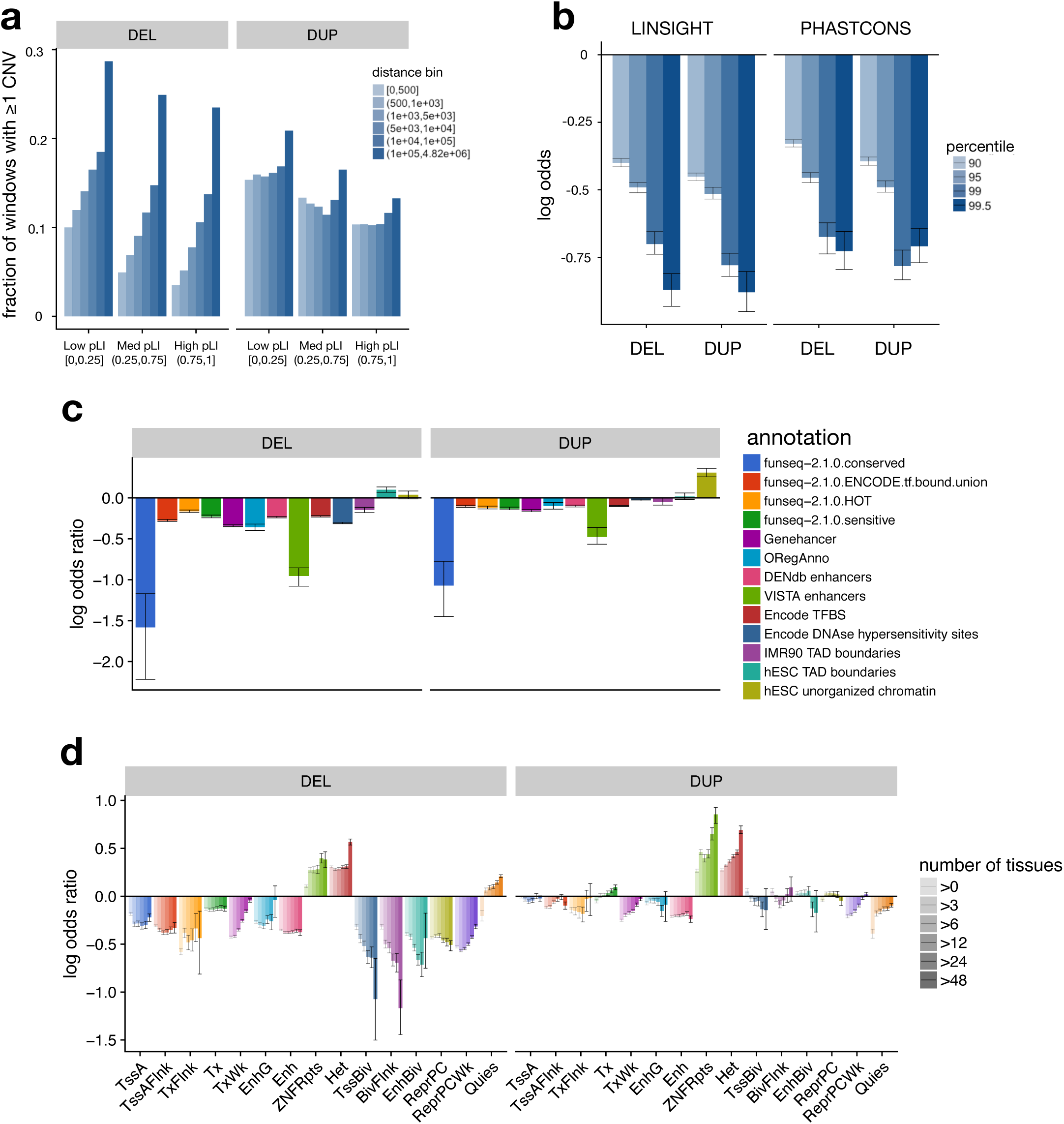
Dosage-sensitivity of functional annotations. **(a)** Fraction of 1 kb genomic windows containing at least one CNV, as a function of distance to the nearest coding exon and the pLI of that gene. **(b)** Depletion of CNV in conserved genomic regions. Odds ratios for the occurrence of CNV in highly conserved (based of LINSIGHT or PHASTCONS percentile) vs. less-conserved regions. Odds ratios estimated by the Cochran-Mantel-Haenszel method and stratified by distance to and pLI of nearest coding exon. Error bars indicate 95% confidence intervals estimated by block bootstrap. **(c)** Log-odds ratios for the occurrence of CNV in 1 kb windows intersecting various functional annotation tracks. **(d)** Log-odds ratios for the occurrence of CNV in 1 kb windows overlapping roadmap segmentations, stratified by the number of roadmap tissues in which the region is observed.

We examined the relative dosage sensitivity of regulatory and epigenomic annotations from various projects^45–50^ (Figs. 5c and 5d). Regulatory elements such as enhancers, polycomb repressors, CTCF sites, DNase hypersensitivity sites, and transcription factor binding sites show strong sensitivity to dosage loss via deletion, whereas inert non-coding annotations such as quiescent and heterochromatic regions do not. The patterns of sensitivity to dosage gains via duplication are broadly similar across annotation classes, albeit weaker. This suggests that, as for genes^44^ (Fig. 5a), dosage sensitivity of non-coding elements is generally consistent with regards to losses versus gains, and does not show obviously distinct patterns at (for example) enhancers, repressors or insulators. Dosage sensitivity of regulatory elements at "bivalent" genomic regions from ROADMAP is greater than their typical counterparts (e.g., enhancers vs. bivalent enhancers in Fig. 5d), suggesting that such elements are under especially strong selection. Interestingly, when we consider the full set of ROADMAP annotations across all 127 cell-types, dosage sensitivity increases gradually as a function of the number of cell-types sharing an annotation at that genomic interval. This suggests that constitutive regulatory elements are more sensitive to dosage changes than those that act in a more cell-type specific manner.

The above analyses offer a first glimpse at the types of SV-based genome annotation efforts that will be possible with increased sample sizes expected to be available in the near future.

## DISCUSSION

Here, we have conducted the largest-scale WGS-based study of SV in the human population to date. Perhaps most notably, the large sample size and use of deep WGS allowed us to map a very large number of rare SVs at very high genomic resolution – typically resolved to a single base – and estimate the burden of deleterious SV relative to other variant classes, which has not been possible in prior studies. Our data suggest that rare SVs account for 4.0-11.2% of deleterious coding alleles and 17.2% of deleterious alleles genome-wide, which is a remarkably large and outsized contribution considering that SVs comprise merely ~0.1% of variants. Especially noteworthy is the surprisingly large burden of rare, strongly deleterious non-coding deletions apparent in our dataset: we estimate that the typical individual carries ~19 rare non-coding deletions that exhibit levels of purifying selection similar to those for high-confidence LoF variants caused by SNVs or indels (of which there are ~34 per individual). Thus, a relatively large number of rare non-coding deletions with strong adverse fitness effects exist in the human population, and it seems reasonable to expect that these make a proportionally large contribution to human disease. These results argue that comprehensive assessment of coding and non-coding SVs will significantly improve trait-mapping power in human genetics studies.

This study has also created valuable community resources. The SVs site-frequency maps that we have generated and publicly released will enable improved SV interpretation efforts. Indeed, there is immense value to having publicly available site-frequency maps from large populations of individuals, generated systematically from modern sequencing platforms and deep data (≥20x), using open source tools and pipelines that are available to the research community (as exemplified by the impact of ExAC/gnomAD^25,26^).

A limitation of this resource is that the false negative rate is expected to be high in repetitive genomic regions and for repetitive SVs including mobile element insertions (MEIs), short tandem repeats (STRs) and multi-allelic CNVs (mCNVs) due to the inherent limitations of algorithms that rely on relatively unique short-read alignments. Indeed, whereas we have reported a mean of 4,442 SVs per genome, recent long-read analyses suggest that as many as ~27,662 SVs may exist per genome when short tandem repeats (STRs) and other highly repetitive forms are counted (https://www.biorxiv.org/content/early/2018/06/13/193144). A subset of repetitive variants are simply impossible to detect using short-reads; others are identified by our pipeline, but are classified as low-confidence SVs. Although it may not be possible to overcome the inherent limitations of short-read sequencing, the comprehensiveness of this resource could be improved in future work through the application of additional specialized SV discovery algorithms tailored to MEIs, STRs and mCNVs; the challenge will be to implement a more comprehensive approach, *at scale*, while maintaining the high computational efficiency and genomic resolution of our current pipeline. For example, current approaches for mapping mCNVs via read-depth analysis^51^ do not scale beyond ~2,000 deep WGS datasets (unpublished observation).

Finally, we have mined this resource to assess the dosage sensitivity of genes and non-coding elements. At genes, our results are consistent with and complementary to existing results based on exome sequencing and array-based methods. The high resolution nature of our SV map also allowed us to examine non-coding elements, where we observed a strong correlation with measures of nucleotide conservation, purifying selection, regulatory element activity, and cell-type specificity. Although our current sample size is inadequate to support informative genome-wide per-base scores required to assess individual non-coding elements, this will be feasible soon as large-scale WGS data resources from various international programs become available for analysis.

Taken together, this work will be invaluable for helping to guide rare variant interpretation in WGS-based human genetics studies.

## ACKNOWLEDGEMENTS

We thank program staff at the National Human Genome Research Institute (NHGRI) for supporting this effort. This study was funded by NHGRI CCDG awards to Washington University in St. Louis (WU) (UM1 HG008853), Broad Institute of MIT and Harvard (UM1 HG008895), Baylor College of Medicine (UM1 HG008898), and New York Genome Center (UM1 HG008901); an NHGRI Genome Sequencing Program (GSP) Coordinating Center grant to Rutgers (U24 HG008956); and a Burroughs Wellcome Fund Career Award to IMH. Additional data production at WU was funded by a separate NHGRI award (5U54HG003079). We thank Shamil Sunyaev for helpful comments on the manuscript. We gratefully acknowledge all individuals involved in the recruitment of samples analyzed for this study. Thanks to Terri Teshiba for coordinating samples for FINRISK and EUFAM sequencing. Data production for EUFAM was funded by 4R01HL113315-05. The METSIM study was supported by grants to Markku Laakso from the Academy of Finland (No. 321428), the Sigrid Juselius Foundation, the Finnish Foundation for Cardiovascular Research, Kuopio University Hospital, and the Centre of Excellence of Cardiovascular and Metabolic Diseases supported by the Academy of Finland. Data collection for the CEPH Pedigrees was funded by the George S. and Dolores Doré Eccles Foundation and NIH grants GM118335 and GM059290. Study recruitment at WU was funded by the DDRCC (NIDDK P30 DK052574) and the Helmsley Charitable Trust. Study recruitment at Cedars-Sinai was supported by the F. Widjaja Foundation Inflammatory Bowel and Immunobiology Research Institute, NIH/NIDDK grants P01 DK046763 and U01 DK062413, and the Helmsley Charitable Trust. Study recruitment at Intermountain Medical Center was funded by the Dell Loy Hansen Heart Foundation. The Late Onset Alzheimer's Disease Study (LOAD) study was funded by grants to T. Foroud (U24AG021886, U24AG056270, U24AG026395, R01AG041797). The Atherosclerosis Risk in Communities (ARIC) study was funded by NHLBI (HHSN268201700001I, HHSN268201700002I, HHSN268201700003I, HHSN268201700004I HHSN268201700005I). The authors thank the staff and participants of the ARIC study for their important contributions. The Population Architecture Using Genomics and Epidemiology (PAGE) program is funded by NHGRI with co-funding from NIMHD (U01HG007416, U01HG007417, U01HG007397, U01HG007376, and U01HG007419). Samples from the BioMe Biobank were provided by The Charles Bronfman Institute for Personalized Medicine at the Icahn School of Medicine at Mount Sinai. The Hispanic Community Health Study/Study of Latinos was carried out as a collaborative study supported by NHLBI (N01-HC65233, N01-HC65234, N01-HC65235, N01-HC65236, N01-HC65237), with contributions from NIMHD, NIDCD, NIDCR, NIDDK, NINDS and NIH ODS. The MEC study is funded through NCI (R37CA54281,R01 CA63, P01CA33619, U01CA136792, and U01CA98758). For the Stanford Global Reference Panel, individuals from Puno, Peru were provided by Drs. Julie Baker and Carlos Bustamante, with funding from the Burroughs Welcome Fund; individuals from Rapa Nui (Easter Island) were provided by Drs. Karla Sandoval Mendoza and Andres Moreno Estrada with funding from the Charles Rosenkranz Prize for Health Care Research in Developing Countries. The WHI program is funded by NHLBI (HHSN268201100046C, HHSN268201100001C, HHSN268201100002C, HHSN268201100003C, HHSN268201100004C, and HHSN271201100004C). The GALA II study and Esteban G. Burchard are supported by the Sandler Family Foundation, the American Asthma Foundation, the RWJF Amos Medical Faculty Development Program, the Harry Wm. and Diana V. Hind Distinguished Professor in Pharmaceutical Sciences II, NHLBI (R01HL117004, R01HL128439, R01HL135156, X01HL134589), NIEHS (R01ES015794, R21ES24844), NIMHD (P60MD006902, R01MD010443, RL5GM118984) and the Tobacco-Related Disease Research Program (24RT-0025). The authors wish to acknowledge the following GALA II co-investigators for subject recruitment, sample processing and quality control: Celeste Eng, Sandra Salazar, Scott Huntsman, MSc, Donglei Hu, PhD, Angel C.Y. Mak, PhD, Lisa Caine, Shannon Thyne, MD, Harold J. Farber, MD, MSPH, Pedro C. Avila, MD, Denise Serebrisky, MD, William Rodriguez-Cintron, MD, Jose R. Rodriguez-Santana, MD, Rajesh Kumar, MD, Luisa N. Borrell, DDS, PhD, Emerita Brigino-Buenaventura, MD, Adam Davis, MA, MPH, Michael A. LeNoir, MD, Kelley Meade, MD, Saunak Sen, PhD and Fred Lurmann, MS. The authors also wish to thank the staff and participants who contributed to the GALA II study.

## METHODS

### Generation of the “Build 38” (B38) callset

#### Per-sample processing

This callset is derived from 23,559 individuals that were part of the CCDG program as well as 950 Latino samples from the PAGE consortium. All data was produced at one of the four CCDG-funded sequencing centers and aligned to genome build GRCh38 using each individual center’s functionally equivalent pipeline implementation^23^. Per-sample calling was performed on 23,547 samples using LUMPY^21^ (v0.2.13), CNVnator^52^ (v0.3.3) and svtyper^22^ (v0.1.4). We excluded HLA, decoy or alternate contigs and regions of much higher than the expected copy number (>12 mean copies per genome across 409 samples) from SV calling with LUMPY (https://github.com/hall-lab/speedseq/blob/master/annotations/exclude.cnvnator_100bp.GRCh38.20170403.bed).

#### Per-sample QC

We observed an excess of small (400 - 1000 bp) singleton deletions (i.e., present in only a single sample), suggesting a large number of false positives. On further investigation, this excess arose from differences between centers in library insert-size distribution. To reduce the number of false positive small deletions, deletions of ≤1000 bp were eliminated unless they had split read support in at least one sample. Subsequently, per-sample quality control was performed to eliminate outlier samples. We removed 213 samples where variant counts (for any SV type) were >6 median absolute deviations from the median count for that type.

#### Merging and cohort-level re-genotyping

The remaining samples were processed into a single, joint callset using svtools (https://www.biorxiv.org/content/early/2018/12/13/494203, https://github.com/hall-lab/svtools) (v0.3.2), modified to allow for multi-stage merging. The code for this merging is available in a container via DockerHub (https://hub.docker.com/) (ernfrid/svtools_merge_beta@sha256:126ad19ad1aae53d05127df93105d83d236ddfb11a8aa65344f0d0aee93 6f919). Samples were merged using svtools lsort followed by svtools lmerge in batches of 1000 samples (or fewer) within each cohort. The resulting per-cohort batches were then merged again using svtools lsort and svtools lmerge to create a single set of variants for the entire set of 23,331 remaining samples. This site list was then used to genotype each candidate site in each sample across the entire cohort using svtyper (v0.1.4). Genotypes for all samples were annotated with copy number information from CNVnator. Subsequently, the per-sample VCFs were combined together using svtools vcfpaste. The resulting VCF was annotated with allele frequencies using svtools afreq, duplicate SVs pruned using svtools prune, variants reclassified using svtools classify (large sample mode), and any identical lines removed. For reclassification of chromosomes X and Y, we used a container hosted on DockerHub (ernfrid/svtools_classifier_fix:v1). All other steps to assemble the cohort above used the same container used for merging.

#### Callset tuning

Using the variant calling control trios, we chose a Mean Sample Quality (MSQ) cutoff for inversions (INV) and breakends (BNDs) that yielded approximately a 5% Mendelian error rate (ME). Inversions passed if: MSQ ≥ 150; neither split-read nor paired-end lumpy evidence made up <10% of total evidence; and support from any one strand was >10%. BNDs passed if MSQ ≥ 250.

#### Genotype refinement

Mobile element insertion (MEI) and deletion (DEL) genotypes were set to missing on a per-sample basis (https://github.com/hall-lab/svtools/blob/develop/scripts/filter_del.py, commit <u>5c32862</u>) if the site was poorly captured by split-reads. Genotypes were set to missing if the size of the DEL or MEI was smaller than the minimum size discriminated at 95% confidence by svtyper (https://github.com/hall-lab/svtools/blob/develop/scripts/del_pe_resolution.py, commit <u>3fc7275</u>). DEL and MEI genotypes for sites with allele frequency ≥0.01 were refined based on clustering of allele balance and copy number values within the datasets produced by each sequencing center (https://github.com/hall-lab/svtools/blob/develop/scripts/geno_refine_12.py, commit <u>41fdd60</u>). In addition, duplications were re-genotyped with more sensitive parameters to better reflect expected allele balance for simple tandem duplications (https://github.com/ernfrid/regenotype/blob/master/resvtyper.py, commit <u>4fadcc4</u>).

#### Filtering for size

The remaining variants were filtered to meet the size definition of an SV (≥50 bp). The length of intra-chromosomal generic breakends (BNDs) was calculated using vawk (https://github.com/cc2qe/vawk) as the difference between the reported positions of each breakpoint.

#### Large callset sample QC

Of the remaining samples, we evaluated per-sample counts of deletions, duplications, and generic breakends within the low allele frequency (0.1% - 1%) class. Samples with variant counts exceeding 10 median absolute deviations from the mean for any of the 3 separate variant classes were removed. In addition, we removed samples with genotype missingness >2%. These QC filters removed a total of 120 additional samples. Finally, we removed 64 samples that were identified as duplicates or twins in a larger set of data.

### Breakpoint resolution

Breakpoint resolution was calculated using bcftools (v1.3.1) query to create a table of confidence intervals for each variant in the callset, but excluding secondary BNDs. Each breakpoint contains two 95% confidence intervals, one each around the start location and end location. For each breakpoint, the resolution was defined as the average width of these two intervals. Summary statistics were calculated in RStudio (v1.0.143; R version 3.3.3).

### Self-reported ethnicity

Self-reported ethnicity was provided for each sample via the sequencing center and aggregated by the NHGRI Genome Sequencing Program (GSP) coordinating center. For each combination of reported ethnicity and ancestry, we assigned a super-population, continent (based on the cohort), and ethnicity. Samples where ancestry was unknown, but the sample was Hispanic, were assigned to the Americas (AMR) superpopulation. Summarized data are presented in Table 1.

### Sample relatedness

As SNV calls were not yet available for all samples at the time of the analysis, relatedness was estimated using large (>1 kb), high-quality autosomal deletions and mobile element insertions with allele frequency >1%. These were converted to plink format using plink (v1.90b3.38) and then subjected to kinship calculation using KING^53^ (v2.0). The resulting output was parsed to build groups of samples connected through first degree relationships (kinship coefficient > 0.177). Each group was assigned an arbitrary family identifier. Correctness was verified by the successful recapitulation of the 36 complete Coriell trios included as variant calling controls.

### Callset summary metrics

Callset summary metrics were calculated by parsing the VCF files with bcftools (v1.3.1) query to create tables containing information for each variant/sample pairing or variant alone, depending on the metric. Breakdowns of the BND class of variation were performed using vawk to calculate orientation classes and sizes. These were summarized using Perl and then transformed and plotted using RStudio (v1.0.143; R version 3.3.3).

### Ultra-rare variant analysis

We defined an ultra-rare variant as any variant unique to one individual or one family of first degree relatives. We expect the false positive rate of ultra-rare variants to be low because systematic false positives due to alignment issues are likely to be observed in multiple unrelated individuals. Therefore, we considered both high and low confidence variants in all ultra-rare analyses.

#### Constructing variant chains

Complex variants were identified as in Chiang et al.^3^ by converting each ultra-rare SV to BED format and, within a given family, clustering breakpoints occurring within 100,000 bp of each other using bedtools^54^ (v2.23.0) cluster. Any clusters linked together by BND variants were merged together. The subsequent collection of variant clusters and linked variant clusters (hereafter referred to as *chains*) were used for both retrogene and complex variant analyses.

#### Manual review

Manual review of variants was performed using IGV (v2.4.0). Variants were converted to BED12 using svtools (v0.3.2) for display within IGV. For each sample, we generated copy number profiles using CNVnator (v0.3.3) in 100 bp windows across all regions contained in the variant chains.

#### Retrogene insertions

Retrogene insertions were identified by examining the ultra-rare variant chains constructed as described above. For each chain, we identified any constituent SV with a reciprocal overlap of 90% to an intron using bedtools (v2.23.0). For each variant chain, the chain was deemed a retrogene insertion if it contained one or more BND variants with +/-strand orientation that overlapped an intron. Additionally, we flagged any chains that contained non-BND SV calls, as their presence was indicative of a potential misclassification, and manually inspected them to determine if they represented a true retrogene insertion.

#### Complex variants

We retained any cluster(s) incorporating 3 or more SV breakpoint calls, but removed SVs identified as retrogene insertions either during manual review or algorithmically. In addition, we excluded one call deemed to be a large, simple variant after manual review.

#### Large variants

Ultra-rare variants >1 Mb in length were selected and any overlap with identified complex variants identified and manually reviewed. Of 5 potential complex variants, one was judged to be a simple variant and included as a simple variant while the rest were clearly complex variants and excluded. Gene overlap was determined as an overlap ≥1 bp with any exon occurring within protein-coding transcripts from Gencode v27 marked as a principal isoform according to APPRIS^55^.

#### Balanced Translocations

Ultra-rare generic “breakend” (BND) variants, of any confidence class, connecting two chromosomes and with support (>10%) from both strand orientations were initially considered as candidate translocations. We further filtered these candidates to require exactly two reported strand orientations indicating reciprocal breakpoints (i.e. +-/-+, −+/+-, --/++, ++/--), no read support from any sample with a homozygous reference genotype, at least one split-read supporting the translocation from samples containing the variant, and <25% overlap of either breakpoint with any simple repeat (downloaded from ftp://hgdownload.cse.ucsc.edu/goldenPath/hg38/database/simpleRepeat.txt.gz).

Comprehensive annotations from the Gencode v27 GTF (ftp://ftp.ebi.ac.uk/pub/databases/gencode/Gencode_human/release_27/gencode.v27.annotation.gtf.gz) were used to determine the number of affected genes. A BED file of all introns was created by converting transcripts and exons to BED entries and subtracting all exons from their respective transcripts using bedtools (v2.23.0). To identify translocations affecting genes, the translocations were converted to BEDPE using svtools (v0.3.1), padded by 1 bp and intersected with introns using bedtools (v2.23.0). The number of unique chromosome/gene name pairs for each translocation was used to determine the number of affected genes affected by each breakpoint.

To determine if a translocation resulted in an in-frame fusion, we converted to BEDPE, padded by 1 bp and intersected the breakpoints with all introns using bedtools (v2.23.0). Each intron entry was then padded by 1 bp and intersected with the Gencode GTF file using bedtools (v2.23.0) and restricting to coding exons of the same transcript as the intron. Then, for each set of exons intersected by a given translocation, all combinations of transcripts were compared, taking into account their orientation and the orientation of the breakpoint, to determine if frame was maintained across the potentially fused exons. The resulting two candidate translocations were manually reviewed by reconstructing the transcript sequence of the fusion and translating the resulting DNA sequence using https://web.expasy.org/translate/ to confirm a single open reading frame was maintained.

### Generation of the "Build 37" (B37) callset

#### Per-sample processing

This callset was constructed starting from a set of 8,455 individuals: 8,181 samples from 8 cohorts sequenced at the McDonnell Genome Institute, as well as 274 samples from the Simons Genome Diversity Project downloaded from EMBL-EBI (https://www.ebi.ac.uk/ena/data/view/PRJEB9586). All samples passed standard production QC metrics and had a mean depth of coverage > 20X. Data were aligned to GRCh37 using the speedseq (v0.1.2) realignment pipeline. Per-sample SV calling was performed with speedseq sv (v0.1.2) using LUMPY (v0.2.11), cnvnator-multi, and svtyper (v0.1.4) on our local compute cluster. For LUMPY SV calling, we excluded high copy number outlier regions derived from >3,000 Finnish samples as described previously (https://www.biorxiv.org/content/early/2018/12/13/494203); https://github.com/hall-lab/speedseq/blob/master/annotations/exclude.cnvnator_100bp.112015.bed).

#### Per-sample QC

Following a summary of per-sample counts, samples with counts of any variant class (DEL, DUP, INV, or BND) exceeding the median plus 10 times the median absolute deviation for that class were excluded from further analysis; 17 such samples were removed.

#### Merging

The remaining samples were processed into a single, joint callset using svtools (v0.3.2) and the two-stage merging workflow (as described above): each of the 9 cohorts was sorted and merged separately in the first stage, and the merged calls from each cohort sorted and merged together in the second stage.

#### Cohort-level re-genotyping

The resulting SV loci were then re-genotyped with svtyper (v0.1.4) and copy-number annotated using svtools (v0.3.2) in parallel, followed by combination of single-sample VCFs, frequency annotation, and pruning using the standard workflow for svtools (v0.3.2). A second round of re-genotyping with more sensitive parameters to better reflect expected allele balance for simple tandem duplications (https://github.com/ernfrid/regenotype/blob/master/resvtyper.py, commit <u>4fadcc4</u>) was then performed, followed by another round of frequency annotation, pruning, and finally reclassification using svtools (v0.3.2) and the standard workflow.

#### Callset tuning and site-level filtering

Genotype calls for samples in 452 self-reported trios were extracted, and Mendelian error rates calculated using a custom R script; we counted as a Mendelian error any child genotype inconsistent with inheritance of exactly one allele from the mother and exactly one allele from the father. Filtering was performed as described for the B38 callset: Inversions passed if: MSQ ≥ 150; neither split-read nor paired-end lumpy evidence made up < 10% of total evidence; and support from any one strand was > 10%. Generic breakends passed if MSQ ≥ 250. SV of length <50 bp were removed, according to our working definition of ‘structural variation’.

#### Final sample-level filtering

Nine samples with retracted consents, and two hydatidiform mole samples were removed from the callset. Subsequently, the numbers of qc-passing, very rare (<0.1% MAF) DEL, DUP, and BND per sample were determined. Excluding the samples in the Simons Genome Diversity cohort (which were expected, in general, to have unusually high counts of rare variants), we determined the median and median absolute deviation (MAD) of the per-sample counts of each type, and excluded outlier samples with a count exceeding the median+10*MAD of any type. Nine samples were removed in this way. Finally, kinship was estimated using KING (v2.0) based on high-quality, autosomal deletion calls, and each SV was annotated in the VCF according to the number of distinct, first-degree family clusters in which it was observed, as for the Build38 callset.

### Build38 SNV/indel callset generation and QC

Per-sample calling was performed at the Broad Institute as part of CCDG joint-calling of 22,609 samples using GATK^56^ (https://www.biorxiv.org/content/early/2018/07/24/201178) HaplotypeCaller v3.5-0-g36282e4. All samples were joint called at the Broad using GATK v4.beta.6, filtered for sites with an excess heterozygosity value of more than 54.69, and recalibrated using VariantRecalibrator with the following features: QD, MQRankSum, ReadPosRankSum, FS, MQ, SOR, and DP. Individual cohorts were subset out of the whole-CCDG callset using Hail v0.2 (https://github.com/hail-is/hail). Following SNV and indel variant recalibration, multiallelic variants were decomposed, and normalized with vt (v0.5)^57^. Duplicate variants and variants with symbolic alleles were then removed. Afterwards, variants were annotated with custom computed allele balance statistics, 1000 Genomes allele frequencies^26^, gnomAD based population data^25^, VEP (v88)^58^, CADD^27^ (v1.2), and LINSIGHT^28^. Variants having greater than 2% missingness were soft filtered. Samples with high rates of missingness (>2%) or with mismatches between reported and genetically-estimated sex (determined using plink v1.90b3.45 sex-check) were excluded. The LOFTEE plugin (v0.2.2-beta; https://github.com/konradjk/loftee) was used to classify putative LoF SNV and indels as high or low confidence.

### Annotation of gene-altering SV calls

The VCF was converted to BEDPE format using svtools vcftobedpe The resulting BEDPE file was intersected (using bedtools (v2.23.0) intersect and pairtobed) with a BED file of coding exons from Gencode v27 with principal transcripts marked according to APPRIS^55^. The following classes of SV were considered putative gene-altering events: (1) DEL, DUP, or MEI intersecting any coding exon; (2) INV intersecting any coding exon and with either breakpoint located within the gene body; and (3) BND with either breakpoint occurring within a coding exon.

### Gene-based estimation of dosage sensitivity

We followed the method of Ruderfer et al.^44^, to estimate genic dosage sensitivity scores using counts of exon-altering deletions and duplications in a combined callset comprising the 14,623 sample pan-CCDG callset plus 3,172 non-redundant samples from the B37 callset. Build37 CNV calls were lifted over to build38 as BED intervals using crossmap (v0.2.1)^59^. We determined the counts of deletions and duplications intersecting coding exons of principal transcripts of any autosomal gene. In Ruderfer et al.^44^, the expected number of CNVs per gene was modeled as a function of several genomic features (GC content, mean read depth, etc.), some of which were relevant to their exome read-depth CNV callset but not to our WGS-based breakpoint mapping lumpy/svtools callset. In order to select the relevant features for prediction, we restricted to the set of genes in which fewer than 1% of samples carried an exon-altering CNV, and used l^1^-regularized logistic regression (from the R glmnet package^60^, v2.0-13), with the penalty λ chosen by 10-fold cross-validation. The selected parameters (gene length, number of targets, and segmental duplications) were then used as covariates in a logistic regression-based calculation of per-gene intolerance to DEL and DUP, similar to that described in Ruderfer et al.^44^. For deletions (or duplications, respectively), we restricted to the set of genes with <1% of samples carrying a DEL, to estimate the parameters of the logistic model. We then applied the fitted model to the full set of genes to calculate genic CNV intolerance scores as the residuals of the logistic regression of CNV frequency on the genomic features, standardized as z-scores and with winsorization of the lower 5^th^ percentile.

### Genome-wide estimation of deleterious variants

In order to estimate the relative numbers of deleterious SNV, indels, DELs and DUPs genome-wide in the normal population, we relied on a subset of 4,298 samples from the B38 callset for which we had joint variant callsets for both SNVs/indels (GATK) and SVs (lumpy/svtools). Each SNV and indel was annotated with CADD^27^ and LINSIGHT^28^ scores as described above. CADD and LINSIGHT scores were converted to percentiles and singleton rates calculated for variants above each score threshold. CADD and LINSIGHT scores were then calibrated to a standard scale by matching singleton rates. Each DEL and DUP was annotated with CADD and LINSIGHT scores, calculated as the mean of the top 10 single-base CADD or LINSIGHT scores, respectively, for the span of the CNV (as implemented in SVSCORE^29^). The CNV-level CADD and LINSIGHT scores were then standardized using the above calibration curves. Finally, each variant (SNV, indel, or CNV) was assigned a combined CADD-LINSIGHT score, calculated as the maximum of the 2 distinct scores.

The combined scores provided a means to rank, within each variant class, variants in order of deleteriousness. We calculated a singleton rate for the set of all LOFTEE high confidence protein-truncating SNV and indels in highly-conserved (top 10% of pLI) autosomal genes. We then estimated the number of deleterious variants of each type genome-wide by choosing the combined CADD-LINSIGHT score threshold as the minimum value such that the singleton rate for the set of higher-scoring variants was greater than or equal to the singleton-rate for LOFTEE high-confidence PTVs.

### Annotation of non-coding elements

We divided the genome into 1 kb non-overlapping windows to investigate the rates of CNV occurrence relative to various classes of coding and non-coding elements, genome-wide. Windows intersecting assembly gaps or high-copy number outlier regions (as described above) and windows with fewer than 50% of bases uniquely mappable as determined using GEM-mappability (build 1.315)^61^ were excluded from analysis. Bed tracks of genomic annotations for the non-coding dosage sensitivity analysis were created as described below.

The phastcons-20way^62^ conservation track was downloaded from the UCSC genome browser (rsync://hgdownload.cse.ucsc.edu/goldenPath/hg38/phastCons20way/hg38.phastCons20way.wigFix.gz) and converted to bed format. The mean phastcons score for each 1 kb window was calculated using bedtools map. Quantiles of mean window-level phastcons scores were calculated and used as thresholds for the sensitivity analysis.

The LINSIGHT^28^ score track was downloaded from CSHL (http://compgen.cshl.edu/~yihuang/tracks/LINSIGHT.bw). The 1kb genomic windows were lifted over to hg19 using crossmap (v0.2.1), annotated with mean per-window LINSIGHT scores using bedtools map and lifted back to GRChb38. Quantiles of mean window-level LINSIGHT scores were calculated and used as thresholds for the sensitivity analysis.

Genehancer^49^ enhancers were downloaded from GeneCards (https://genecards.weizmann.ac.il/geneloc/index.shtml) and converted to bed format.

Vista^48^ enhancers were downloaded from LBL (https://enhancer.lbl.gov/cgi-bin/imagedb3.pl?page_size=20000;show=1;search.result=yes;page=1;form=search;search.form=no;action=search;search.sequence=1), restricted to human enhancers, converted to bed format and lifted over to GRChb38 using crossmap.

Encode^45^ DNAse hypersensitivity sites and transcription factor binding sites were downloaded from UCSC (http://hgdownload.cse.ucsc.edu/goldenPath/hg19/encodeDCC/wgEncodeRegDnaseClustered/wgEncodeRegDnaseClusteredV3.bed.gz, http://hgdownload.cse.ucsc.edu/goldenPath/hg19/encodeDCC/wgEncodeRegTfbsClustered/wgEncodeRegTfbsClusteredV3.bed.gz) and lifted over to GRCHb38 using crossmap.

Oreganno^63^ literature-curated enhancers were downloaded from UCSC (http://hgdownload.cse.ucsc.edu/goldenpath/hg19/database/oreganno.txt.gz) converted to bed format, and lifted over to GRChb38 using crossmap.

Sensitive^47^, transcription factor bound, ultra-conserved^64^, and HOT^65^ regions were downloaded from the funseq2^66^ resources (http://archive.gersteinlab.org/funseq2.1.0_data).

Dragon enhancers were downloaded from DENdb^67^ (http://www.cbrc.kaust.edu.sa/dendb/src/enhancers.csv.zip), converted to bed format, lifted over to GRChb37, and filtered for score>2.

Chromatin interaction domains derived from Hi-C on hESC and IMR90 cells^68^ were downloaded from http://compbio.med.harvard.edu/modencode/webpage/hic/, and distances between adjacent topological domains calculated with bedtools. When the physical distance between adjacent topological domains was <400 kb, these were classified as TAD boundaries; otherwise, they were classified as unorganized chromatin. The TAD boundaries and unorganized chromatin data were converted to bed format and lifted over to GRCh38 using crossmap.

Roadmap chromatin state segmentations for 127 epigenomes were downloaded from Roadmap^46^ (https://egg2.wustl.edu/roadmap/data/byFileType/chromhmmSegmentations/ChmmModels/coreMarks/jointModel/final/) and lifted over to GRCh38. Bedtools multiinter was used to determine the number of epigenomes in which each segment was present.

### Dosage sensitivity of non-coding elements

To maximize power, DEL and DUP calls from the non-redundant union of the B37 and B38 callsets (as described above) were used for this analysis. Each window was further characterized by its distance to the nearest exon (the minimum distance between any point in the window and any point in the exon) and the pLI score of the gene corresponding to the nearest exon. The pLI score was set to zero for genes with pLI undefined. In the event that exons of 2 genes were equidistant to the window, the max of the two pLI scores was selected.

For a given SV type (DUP or DEL) and a given functional annotation (e.g., VISTA enhancers), each window was characterized by the presence or absence of one or more SV and the presence or absence of one of more genomic features. We observed a depletion of CNV in windows near exons, and in particular near exons of LoF-intolerant genes (see Fig. 5a). As such, we used a Cochran-Mantel-Haenszel test to estimate the odds ratios for each SV type/functional annotation, while stratifying for the proximity to the nearest exon as well as that exon’s LOF-intolerance (pLI). Because adjacent windows are not strictly independent observations – i.e., CNV or features may overlap adjacent windows inducing some spatial correlations – we used a block bootstrap method (resampling was performed on non-overlapping blocks of 10 windows) to estimate robust confidence intervals.

---

**Supplementary Figure 1.**
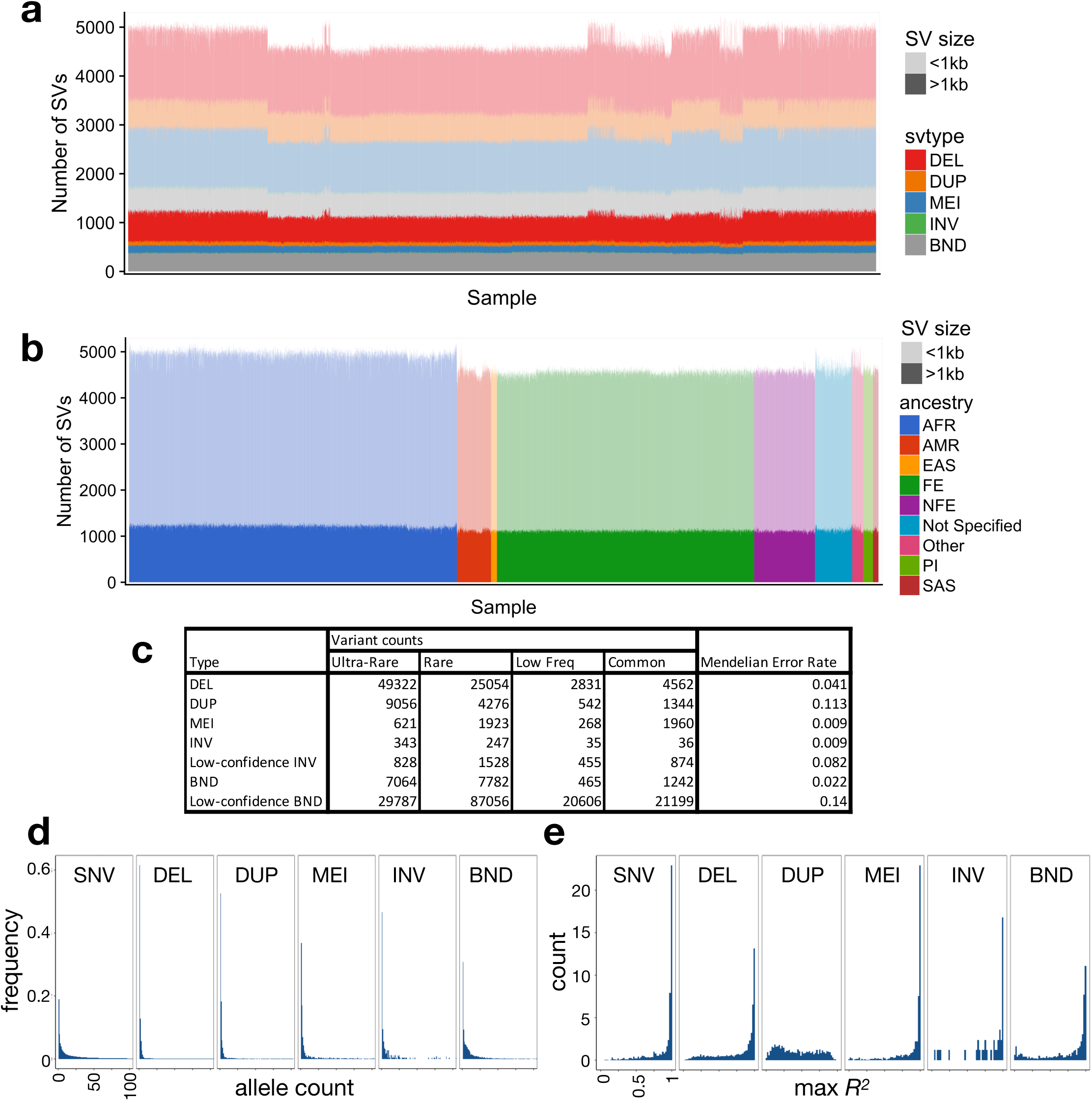
The B37 callset. **(a)** Variant counts (y-axis) for each sample (x-axis) in the callset, ordered by cohort, where large (>1 kb) variants are shown in dark shades and smaller variants in light shades. **(b)** Variant counts per sample, where samples are ordered by self-reported ancestry according to the color scheme at right, using the abbreviations described in Table 1. Note that African-ancestry samples show more variant calls, as expected. **(c)** Table showing the number of variant calls by variant and frequency class, and Mendelian error rate by variant type. **(d)** Histogram of allele count for each variant class, showing alleles with counts ≤100. **(e)** Linkage disequilibrium of each variant class as represented by max *R2* value to nearby SNVs. Note that these distributions mirror those from our prior SV callset for GTEx^3^, which was characterized extensively in the context of eQTLs.

**Supplementary Figure 2.**
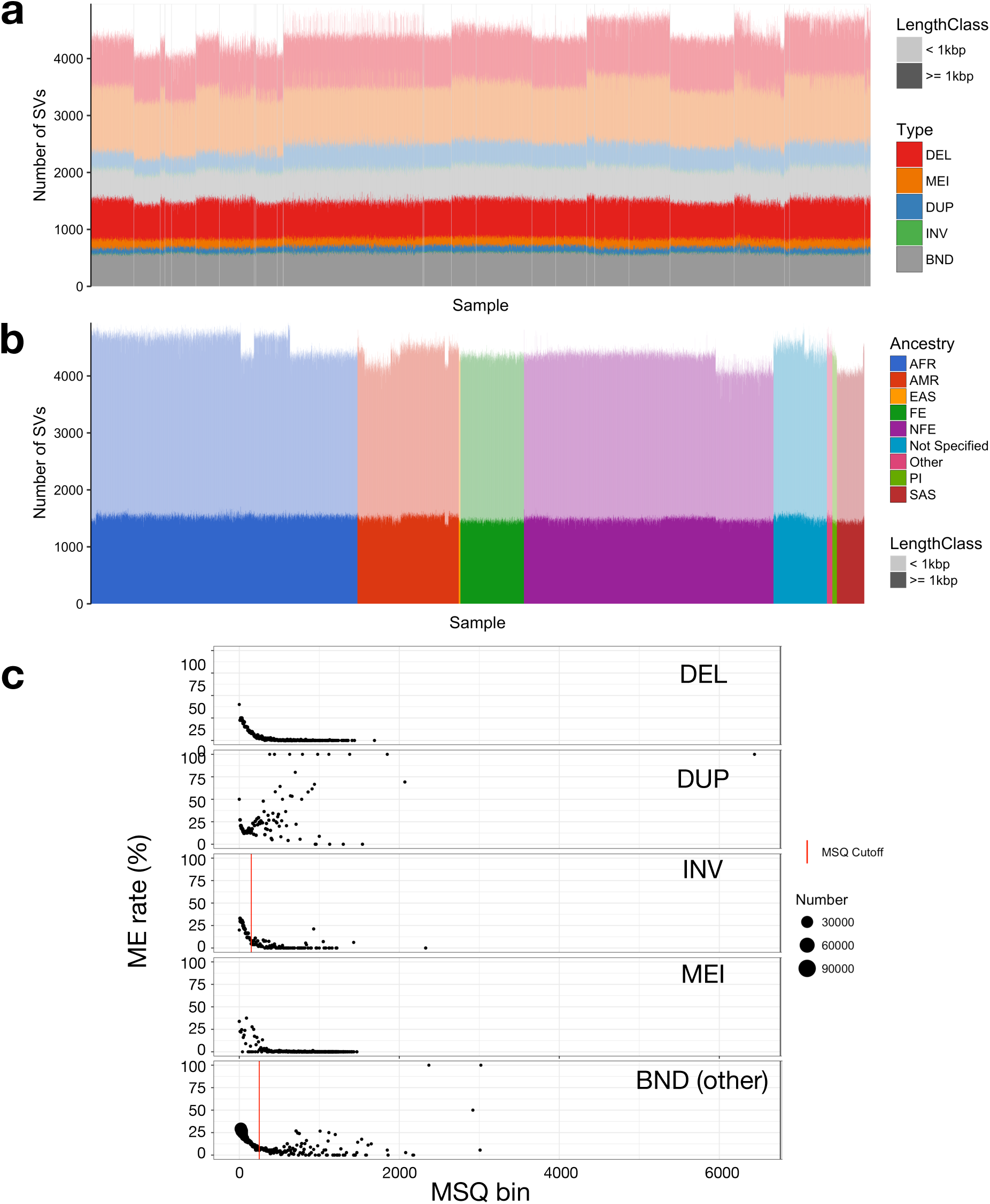
The B38 callset. **(a)** Variant counts (y-axis) for each sample (x-axis) in the callset, ordered by cohort (separated by vertical lines), where large (>1 kb) variants are shown in dark shades and smaller variants in light shades. **(b)** Variant counts per sample, where samples are ordered by self-reported ancestry according to the color scheme at right, using the abbreviations described in Table 1. Note that African-ancestry samples show more variant calls, as expected. Note also that there is some residual variability in variant counts due to differences in data from each sequencing center, but that this is limited to small tandem duplications (see part a), primarily at STRs. **(c)** Plots of Mendelian error (ME) rate (y-axis) by mean sample quality (MSQ) for each variant class, where dot size is determined by point density (see right) and the threshold used to determine high and low confidence SVs is shown by the vertical lines.

**Supplementary Figure 3.**
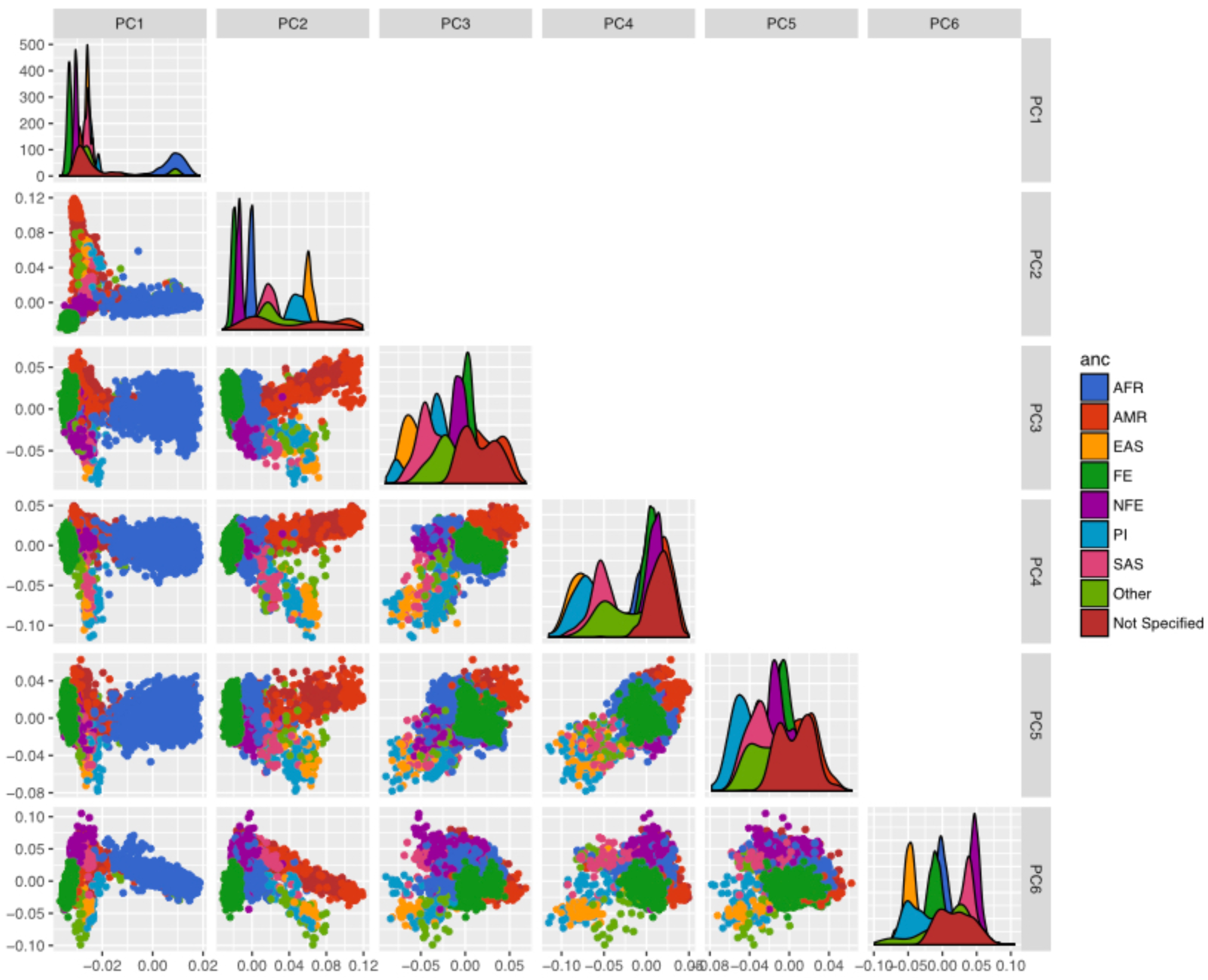
Principal components analysis for the B37 callset. PCA were calculated using an LD-pruned subset of high-confidence DEL and MEI variants, with MAF>1%. Ancestry is based on self-report, using the color scheme at right, using the ancestry abbreviations described in Table 1.

**Supplementary Figure 4.**
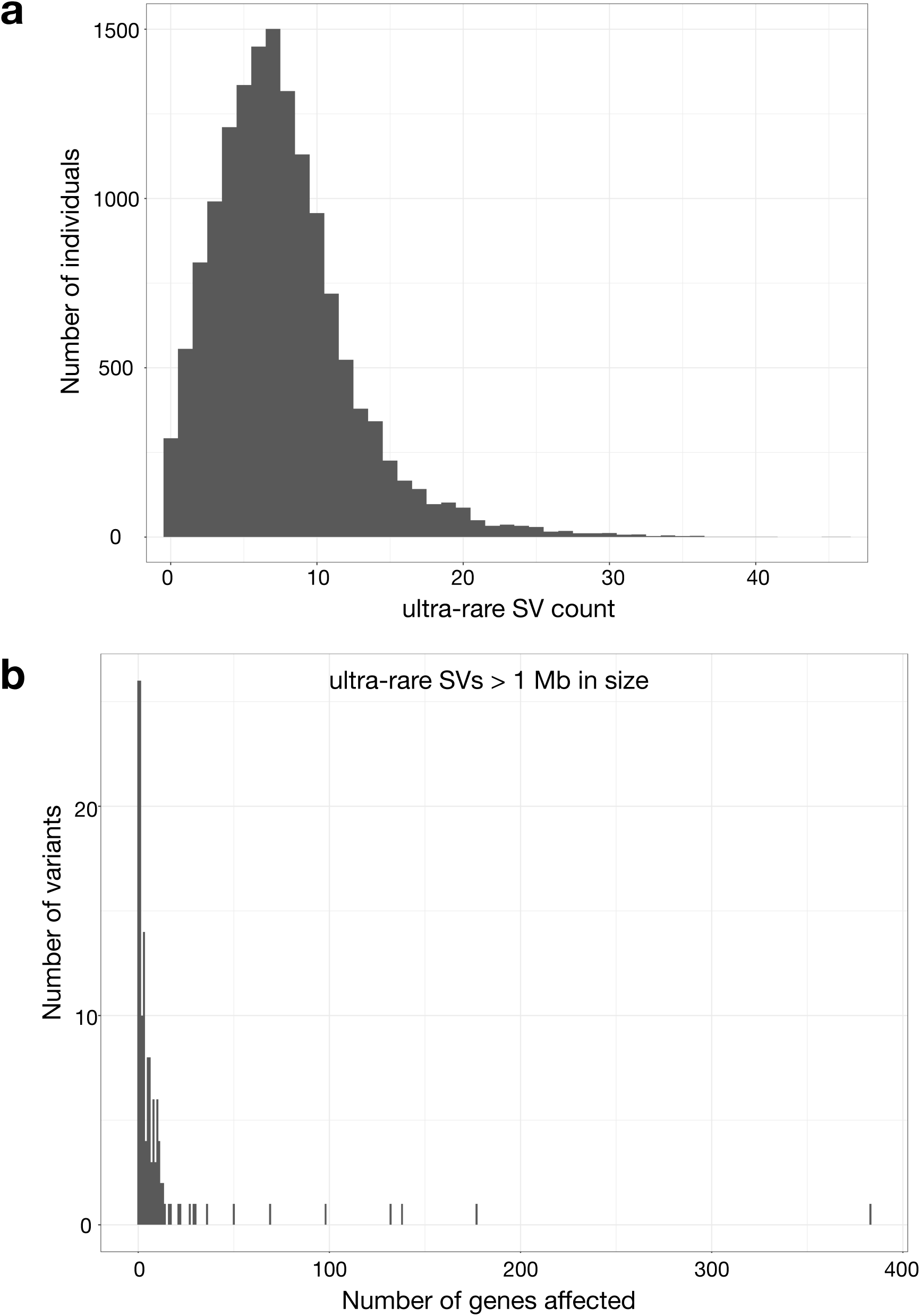
Ultra-rare SVs. **(a)** Histogram showing the number of ultra-rare SVs per individual, where ultra-rare is defined as "singleton" variants private to single individual or nuclear family. **(b)** Histogram showing the number of genes affect by ultra-rare SVs larger than 1 Mb in size.

**Supplementary Figure 5.**
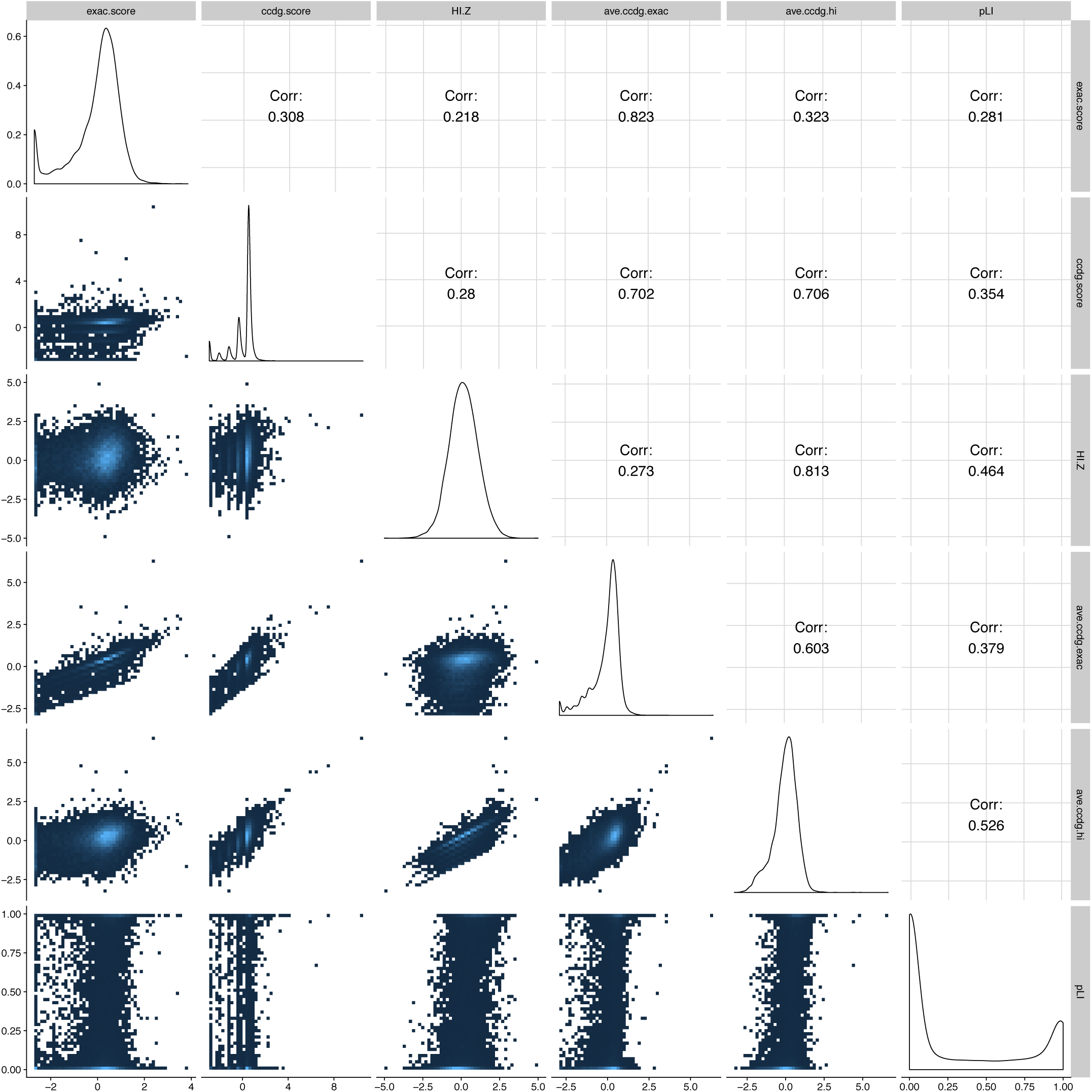
Correlations between dosage sensitivity scores for deletions. ExAC score is the published ExAC DEL intolerance score. CCDG score is similarly calculated, using CCDG deletions. pLI is the published pLI score, HI.Z is the negative of the inverse-normal transformed DECIPHER HI score. Ave.ccdg.exac is the arithmetic mean of the CCDG and ExAC scores. Ave.ccdg.hi is the arithmetic mean of the CCDG and HI-Z scores. Correlations shown are Spearman rank correlations.

**Supplementary Figure 6.**
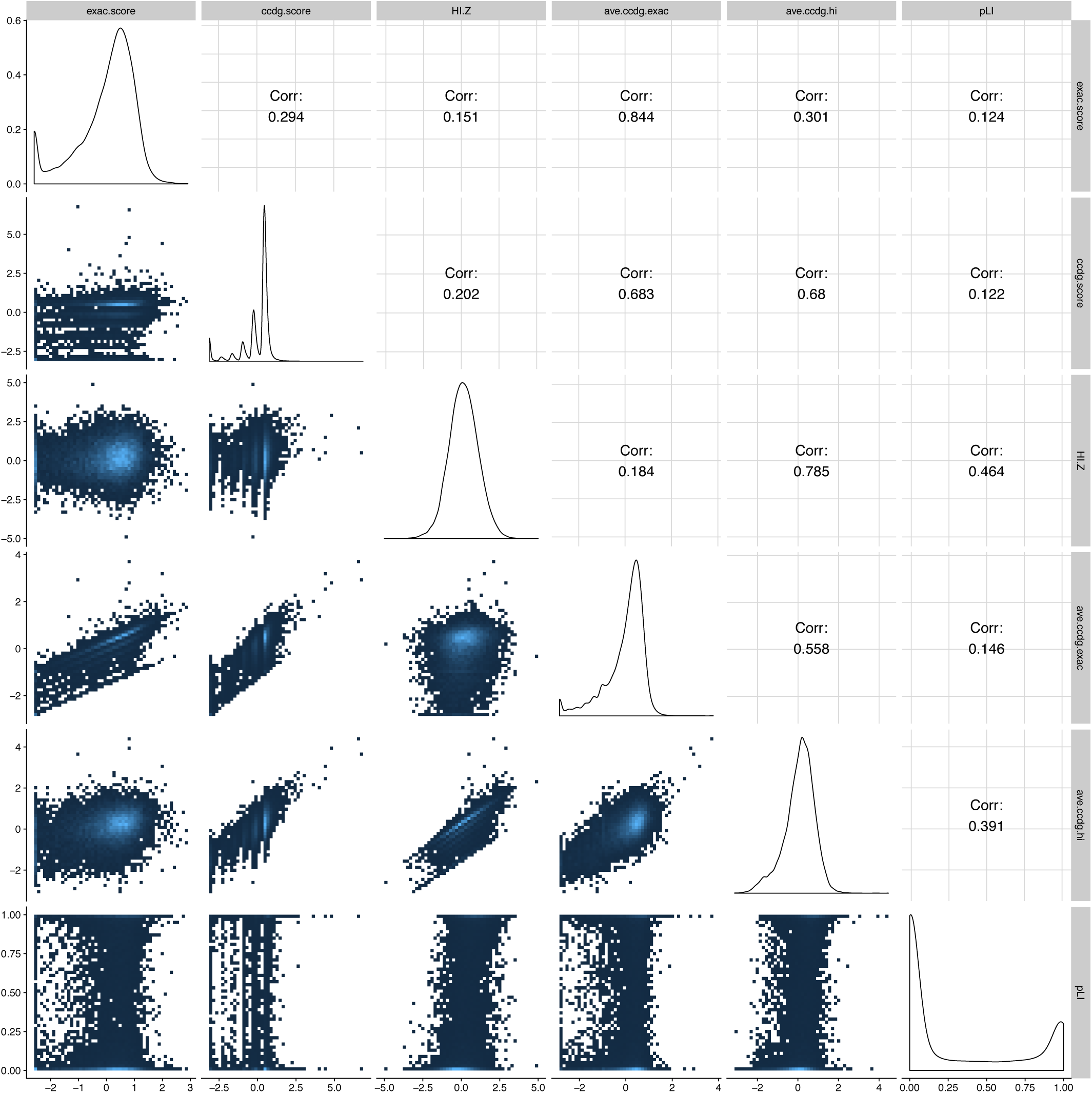
Correlations between dosage sensitivity scores for duplications. ExAC score is the published ExAC DUP intolerance score. CCDG score is similarly calculated, using CCDG duplications. pLI is the published pLI score, HI.Z is the negative of the inverse-normal transformed DECIPHER HI score. Ave.ccdg.exac is the arithmetic mean of the CCDG and ExAC scores. Ave.ccdg.hi is the arithmetic mean of the CCDG and HI-Z scores. Correlations shown are Spearman rank correlations.

